# Canonical self-supervised pretraining paradigm constrains the capacity of genomic language models on regulatory decoding

**DOI:** 10.64898/2026.04.13.715198

**Authors:** Ye-Xi Liang, Yu Wang, Wang-Yue Pan, Zi-Yu Chen, Jia-Chen Wei, Ge Gao

**Affiliations:** State Key Laboratory of Gene Function and Modulation Research, School of Life Sciences, Biomedical Pioneering Innovative Center (BIOPIC) & Beijing Advanced Innovation Center for Genomics (ICG), Center for Bioinformatics (CBI), Peking University, Beijing 100871, China; Changping Laboratory, Yard 28, Science Park Road, Changping District, Beijing 102206, China; Academy for Advanced Interdisciplinary Studies, Peking University, Beijing 100871, China

**Author notes:** These authors contribute equally (Ye-Xi Liang, Yu Wang). Corresponding authors. (Yu Wang), (Ge Gao).

**Keywords:** Genomic language model, Benchmark, Pretraining objective, Gene regulation, Functional genomics

## Abstract

Recent studies suggest that genomic language models (gLMs) could help decode genomic regulatory code. Here, we systematically evaluated 11 representative gLMs across multiple regulatory genomics applications and found that current gLMs offer limited advantages over the random baseline. Further analysis revealed a systematic misalignment between the canonical sequence-only self-supervised pretraining paradigm and the context-specific dynamic nature of gene regulation, highlighting the need for function-oriented pretraining strategies that explicitly incorporate biochemical and regulatory priors.

## Introduction

Large language models have revolutionized the modeling of complex sequential data through self-supervised learning on unlabeled data. Building on the success in modeling human language^1^, researchers are actively resembling such strategy to develop multiple genomic language models (gLM) for learning the regulatory code in human genome^2–12^. Most of these gLMs employ the pretraining-finetuning paradigm: models are pretrained on large-scale unlabeled genomic sequences to predict masked nucleotides (tokens) from their sequence context, and the learned representations are subsequently adapted to downstream applications through task-specific fine-tuning, based on the assumption that the sequence-only self-supervised learning paradigm is sufficient to model the functional regulatory logic encoded in genomic sequences.

However, this assumption remains largely untested. Unlike human language, genomic sequences have different evolutionary constraints, organizational principles, and functional properties, raising questions about whether such sequence-centric paradigms are sufficient for fully capturing the biological meaning of genome regulation.

To bridge such gap, we present LingoDNABench, a comprehensive regulatory-oriented benchmark suite designed to evaluate whether gLMs can extract transferable sequence representations across the full regulatory hierarchy: the prediction of chromatin profiling, transcription regulation, post-transcription regulation, and gene expression. **(Fig. 1a)**. We systematically evaluated 11 representative gLMs spanning diverse model architectures, parameter scales, pretraining data corpora, and pretraining objectives: Caduceus^2^, HyenaDNA^3^, DeepGene^4^, GPN-MSA^5^, DNABERT^6^, DNABERT-2^7^, OmniNA^8^, LucaOne^9^, Nucleotide Transformer^10^, GENERator^11^ and Evo2^12^ (**Methods**). Surprisingly, our finding reveals that these gLMs yield marginal and inconsistent improvements across most applications, even when comparing with random baseline. We further demonstrate that the popular masking-based pretraining paradigm exhibits a fundamental misalignment with most applications, except disease-related variant effect prediction, which is conservation-driven.

**Fig. 1:**
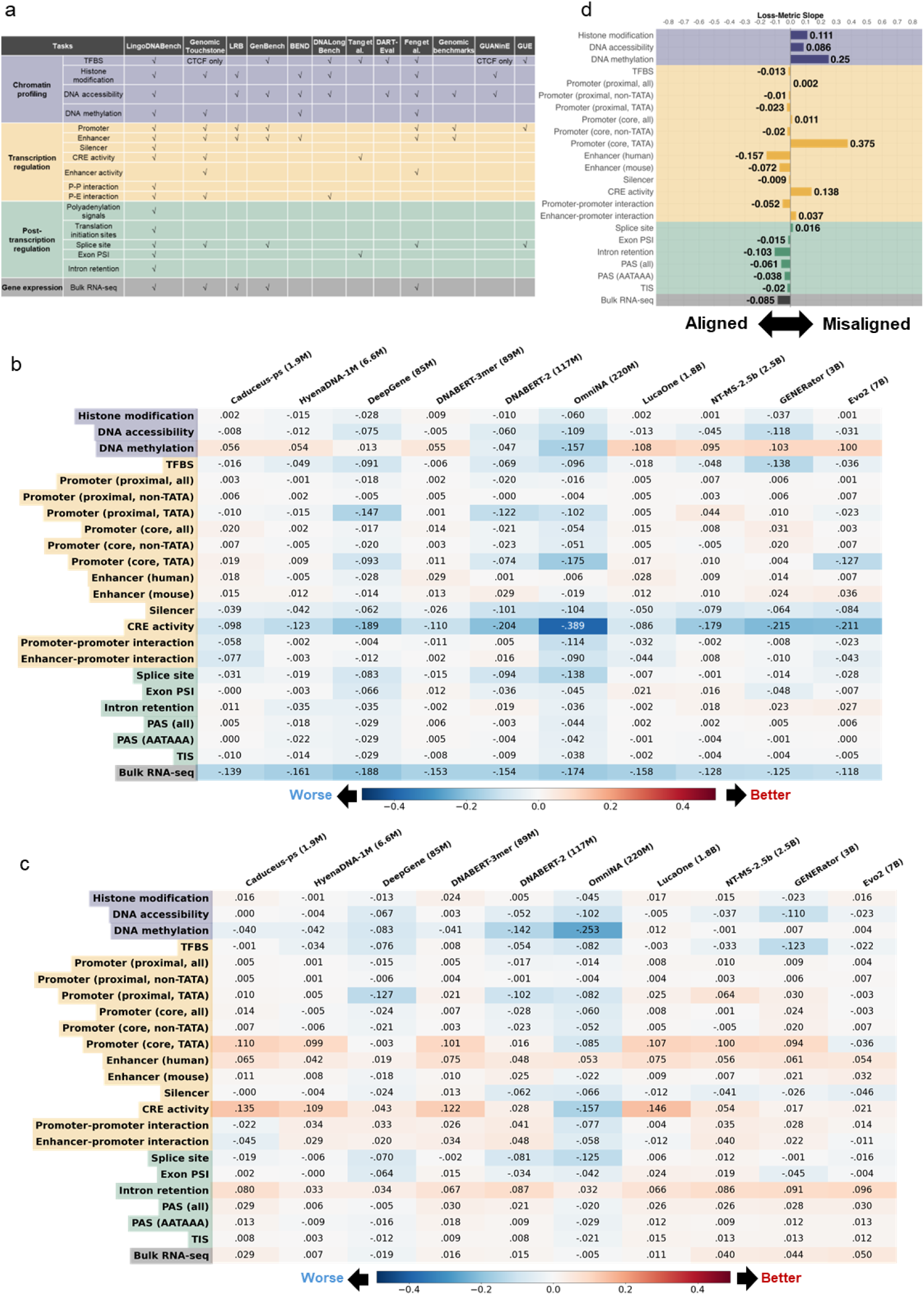
Evaluation of genomic language models on regulatory-oriented downstream applications. **a.** The LingoDNABench evaluation suite, comprises four regulatory categories: chromatin profiling, transcriptional regulation, post-transcriptional regulation and gene expression prediction. **b.** Performance gap between gLMs and non-gLM baselines (*metric*_*gLM*_ −*metric*_*non*−*gLM baselines*_). Red: gLMs superior; blue: baseline superior, same as in (c). The most notable improvement was observed for DNA methylation prediction where 40% gLMs achieving ∼10% enhancement over non-gLM baselines. **c.** Performance gap between gLMs and RandomWeight baseline (*metric*_*gLM*_ − *metric*_*RandomWeight*_). **d.** Alignment between pretraining loss and downstream performance, quantified by the regression slope. More negative slope indicates a stronger alignment.

## Results

A primary objective of gLMs is to extract transferable sequence representations that encode regulatory information across multiple biological scales—from local functional patterns to integrated gene-level outcomes. To evaluate whether gLMs achieve this goal, we developed LingoDNABench, a comprehensive benchmark spanning four regulatory categories across the full regulatory hierarchy: chromatin profiling, transcriptional regulation, post-transcriptional regulation, and gene expression prediction, collectively recapitulating the flow of genetic information (**Methods**).

Specifically, chromatin profiling applications assess the model’s ability to capture local epigenetic states^13^; transcriptional regulation applications evaluate the recognition of *cis*-regulatory elements and their combinatorial interactions, thereby probing higher-order regulatory logic beyond local motifs^14^; post-transcriptional regulation applications determine whether gLM sequence representations encode signals governing RNA processing and translation^15^; gene expression prediction serves as an integrated functional readout, reflecting the culmination of these multi-layered regulatory processes^14^.

To ensure a high-fidelity assessment of sequence representations of pretrained gLMs, we utilized fine-tuning with lightweight adapters. This strategy preserves the pretrained backbone while enabling task-specific adaptation, ensuring that observed downstream performance are directly attributable to the intrinsic information captured during pretraining rather than confounding architectural advantages^16,17^.

We first took non-gLM tools trained specifically for particular task as baselines (**Methods**). No gLM shows a consistent performance advantage over non-gLM baselines among 23 evaluated tasks **(Fig. 1b)**: non-gLM baselines outperformed at least one gLM by more than 5% in 15 out of 23 tasks, surpassing by up to 38.9%.

Inspired by Vishniakov *et al.*^18^ we further introduced RandomWeight model as random baseline, initialized with random parameters and devoid of any training (**Methods**). Surprisingly, all gLMs show marginal improvements for the RandomWeight, suggesting that the current gLMs hardly learn *bona fide* regulatory grammar **(Fig. 1c)**.

One critical assumption for pretraining-based gLMs is that lower pretraining loss could be translated to improved performance for downstream applications^19^. To evaluate this assumption empirically, the longitudinal tracking for both pretraining loss and downstream performance across multiple pretraining epochs is needed. Given the fact that existing gLMs typically do not release intermediate pretraining checkpoints, we pretrained a BERT-based gLM using multiple species genomic sequences (model **M**, **Methods**). Given that the model **M** achieved comparable performance with state-of-the-art gLMs on LingoDNABench (Supplementary Fig. 1), we use it as the representative model for follow-up empirical analysis.

By probing the longitudinal correlation between pretraining loss and downstream performance, we found a better pretraining does not necessarily translate to *bona fide* performance gains of real world regulatory applications like functional element or gene expression prediction applications, which is consistently with the observed inferior performances and suggest a systematic misalignment between the pretraining and downstream objectives (**Fig. 1d** and **Methods**).

We proposed a theoretical framework grounded from an information theory perspective for masking-based pretraining (**Methods**). Briefly, the pretraining objective is essentially minimizing the Kullback-Leibler (KL) between the model’s fitted and the *bona fide* conditional distribution of masked tokens given their sequence context. In other word, this optimization procedure tries to maximize the mutual information between masked tokens and their context, i.e. the model effectively captures the intra-sequence statistical correlation structure. Biologically, the pretraining phase could be considered as a procedure to identify sequences with recurring patterns, i.e. evolutionary conserved regions when training with multiple species and repetitive motifs when training with single specie genome only.

The framework suggests masked-token prediction accuracy should vary across genomic regions according to their intrinsic sequence regularity. As expected, we found that most evaluated gLMs achieve higher masked-token prediction accuracy on elements with strong local sequence correlations (like repetitive elements), while the random baseline performed uniformly (**Fig. 2a** and **Methods**). Consistent with its biology-tuned pretraining strategy including down-weights eukaryotic repetitive elements and incorporating large volume of prokaryotic genomes, the Evo2 showed no significant accuracy disparity between exons and Alu elements.

**Fig. 2:**
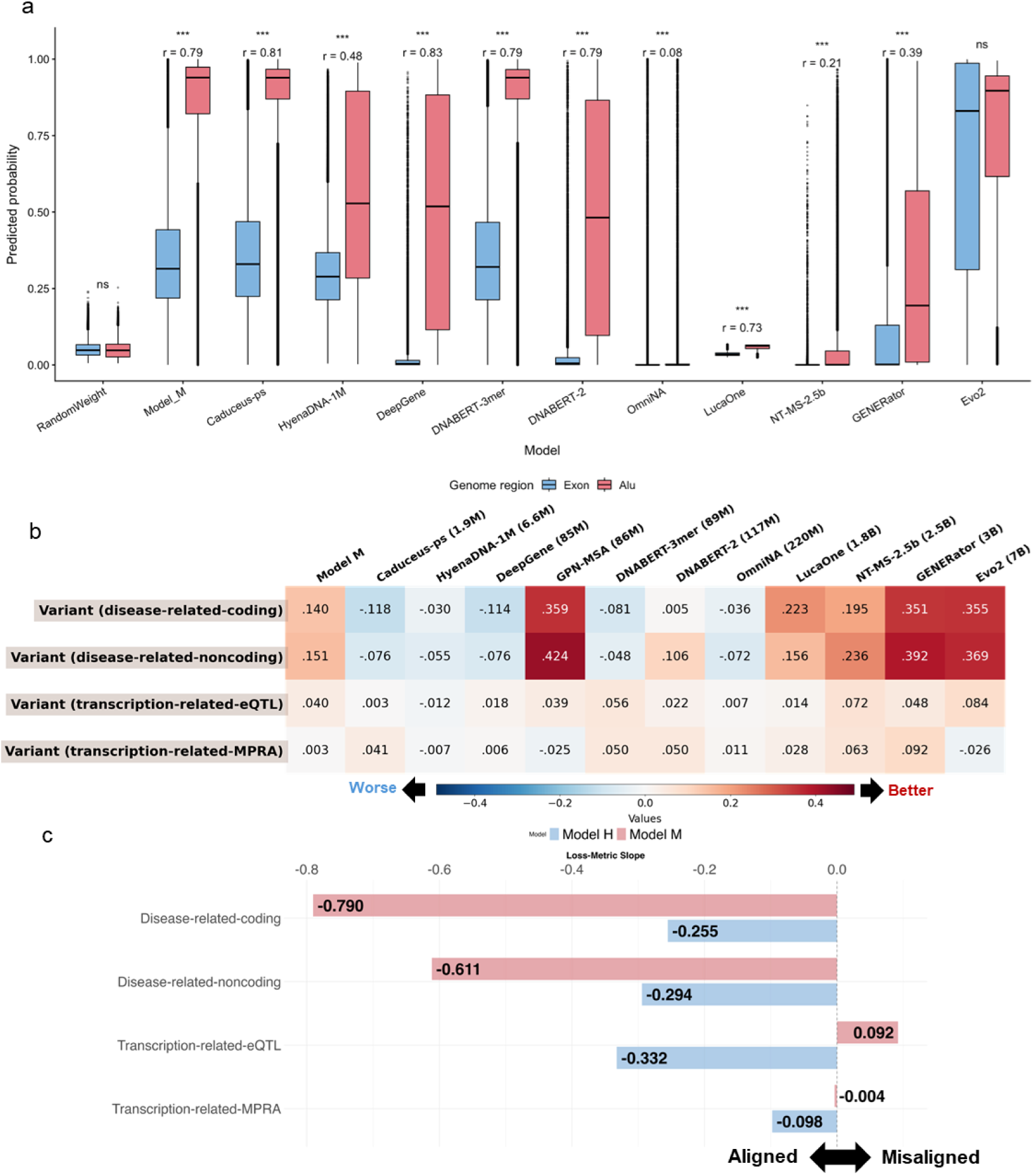
Genomic language models capture intra- and inter-species mutual information via MLM pretraining objective. **a.** Masked-token prediction accuracy across different genomic regions (exons versus Alu elements). Statistical significance was assessed using a one-sided Mann– Whitney U test; *** denotes *P* < 0.001, and “ns” indicates not significant. Effect sizes are reported as rank-biserial correlation (*r*), with positive values indicating higher predicted probabilities in exonic regions relative to Alu elements. In each boxplot, the lower and upper boundaries of the box represent the first (Q1) and third quartiles (Q3), with the median indicated by a line inside the box. The whiskers typically extend to the most extreme data points within 1.5 times the interquartile range (IQR) from the quartiles. Data points outside this range are considered outliers and are plotted individually by points. **b.** The performance of zero-shot variant effect prediction for baselines and gLMs. Disease-related VEP datasets was derived from ClinVar, which are strongly stratified by conservation metrics, and transcription-related VEP datasets with conservation-independent effects served as controls. **c.** Alignment between pretraining loss and performance of variant effect prediction, quantified by the regression slope. **d.** Performance gap between gLMs and RandomWeight baseline n variant effect prediction (metric_gLM-metric_RandomWeight). Red: gLMs superior; blue: baseline superior.

Guided by this framework, we hypothesized that current gLMs are inherently biased toward capturing sequence features shaped by deep evolutionary history. To test this, we designed a comparative evaluation, implementing variant effect prediction evaluation tasks using datasets with differentially discriminable by evolutionary conservation (**Methods**).

We first assembled disease-related genetic variant datasets derived from ClinVar, which are strongly stratified by conservation metrics (Supplementary Table 1). Consistent with our hypothesis, we found most gLMs pretrained on multi-species genomes significantly outperformed random baseline on this variant datasets (**Fig. 2b**), using zero-shot variant prioritization approach (**Methods**), a standard evaluation setting for variant effect prediction^20^.

To determine whether this performance gain stems from the conservation solely, we further compiled eQTL and MPRA-based genetic variant datasets as controls. Unlike disease-related variant datasets, these datasets exhibit minor differences in evolutionary conservation between transcription-modulating and non-transcription-modulating loci (Supplementary Table 1), thereby serving as a test for the detection of biochemically-driven regulatory effects. Under same evaluation protocols, we found that gLM performance dropped precipitously on these regulatory-centric controls (**Fig. 2b**).

To further elucidate whether this evolutionary constrain is explicitly driven by the diversity of the pretraining corpus, we performed a longitudinal analysis of pretraining dynamics across different species scales. To test this, we complemented previously described multi-species model **M** with a human-specific one (model **H**) using identical protocol (**Methods**). On disease-related variants, we found model **M** exhibited a clear negative correlation between pretraining loss and variant prioritization performance, indicating that continued optimization primarily strengthens the encoding of evolutionary constraints. In contrast, model **H** showed only marginal, pretraining loss-independent performance gains, confirming that single-species pretraining is insufficient to recover evolutionary signals. Crucially, no such scaling relationship—where improved pretraining loss translates to higher downstream performance—was observed for transcription-related variants in both models (**Fig. 2c**).

## Discussion

Our study provides a critical interrogation of a central assumption underlying current gLMs: the sequence-only self-supervised pretraining is sufficient to support regulatory decoding. First, we present the most comprehensive regulatory-oriented benchmark for gLMs, LingoDNABench. Then, we systematically evaluated multiple representative gLMs and found limited benefits over various baselines in general. In particular, both theoretical and empirical analyses suggest that, while current sequence-only self-supervised pretraining encodes recurrent/conserved patterns effectively, it offers little advantage for tasks whose key signals come from mechanistic interactions like TF binding and gene expression regulation.

Notably, most contemporary gLM pretraining strategies are directly adapted from natural language processing (NLP), with masked language modeling (MLM) as the dominant objective. By design, MLM optimizes for prediction of masked tokens based on given context, a process that inherently prioritizes distributional regularities and statistical co-occurrences.

This parallels insights from classical NLP theory, where MLM objectives are known to emphasize co-occurrence patterns and distributional regularities, which are insufficient for inducing deeper semantic representations required for complex reasonsing^21,22^. Given the fact that multiple NLP downstream tasks can be reformulated into forms that align with the pretraining objective of masked token prediction^23,24^, the information learned during pretraining can be transferred naturally to a wide range of downstream applications^25,26^.

On the other hand, gene regulatory is highly dynamic process determined by both *cis-* elements and *trans-* factors. The canonical MLM pretraining objective enables gLMs to effectively capture evolutionary conservation signals^5,27^, but not dynamic cellular context^28,29^. Meanwhile, the fact that regulatory elements usually evolve under weaker evolutionary constraint than coding regions^30,31^ further limits its utility in real world. Consistently, we also noticed that a recent analysis reveals that the pretraining loss of protein language models correlates with the performance of structure-prediction only, not for protein-property prediction^19^, in addition to several recent benchmarking studies of gLMs^16,32,33^.

In light of this systematic and fundamental limitation, function-oriented paradigms with biochemical and regulatory priors incorporated explicitly offer a plausible alternative. Models like Enformer^14^, SVEN^34^, and AlphaGenome^35^ demonstrate that supervised training on multi-modal regulatory assays substantially improves chromatin and transcriptional predictions. The recent Nona framework introduced a hybrid pretraining strategy that jointly leverages genomic sequence and functional genomics^36^. Along a similar direction, Nucleotide Transformer v3 combines genome-scale MLM pretraining with a multispecies post-training stage that integrates supervised functional genomics and genome annotation, substantially improving regulatory prediction beyond sequence-only pretraining^37^. However, these approaches require extensive, high-quality functional genomics data, which remain scarce, sparse, and unevenly distributed across cell types and conditions. This data bottlenecks fundamentally restricts generalization, hindering broad regulatory prediction.

Collectively, our findings demonstrate that current gLM pretraining, while proficient at distilling evolutionary history, remains fundamentally unaligned with the functional requirements of regulatory decoding. This suggests that the conventional “scaling law" is insufficient for capturing the non-linear, tissue-specific logic of the human genome. Bridging this gap require a paradigm shift: moving beyond the mere expansion of sequence corpora toward the integration of multi-modal functional priors and the development of architectures capable of explicitly encoding the complex orchestration of *cis*- and *trans-*regulatory dynamics. Future advancements must prioritize the transition from models that recognize statistical recurrence to those that decipher the regulatory grammar governing the functional genome.

## Methods

### Construction of LingoDNABench benchmark suites

#### Chromatin profiling

##### DNA accessibility prediction

We downloaded DNase-seq and ATAC-seq data (narrowPeak files, GRCh38) from ENCODE (version 240317), collecting 1,770 raw files in total. Chromatin features from different cell lines and experimental treatments were treated as distinct features (prediction targets). Chromatin features derived from distinct cell lines or treatment methods were treated as separate prediction targets. Following the Basset pipeline^38^, peaks were first extended symmetrically to 510 bp, and overlapping peaks (>200 bp) were then iteratively merged until convergence. Merged regions were assigned binary labels and converted to FASTA sequences from the positive strand.

The final dataset comprised 1,770 distinct features. Sequences from chromosomes 2 and 4 were considered as validation set (496,363 sequences); chromosomes 1, 8, and 9 as test set (619,584 sequences) and the remaining chromosomes as training set (2,414,452 sequences).

##### Histone modification prediction

We downloaded ChIP-seq data (narrowPeak files, GRCh38) from ENCODE (version 240317), collecting 3,070 raw files. The construction of dataset was identical to DNA accessibility data. The final dataset comprised 3,070 distinct features, with 3,741,461 sequences in the training set, 775,558 in the validation set, and 929,143 in the test set.

##### DNA methylation prediction

We employed two datasets for predicting 5-methylcytosine (5mC) and N6-methyladenosine (6mA) in human from iDNA-MS^39^. Positive samples consisted of 41-bp sequences centered on experimentally validated methylation sites; negative samples were sequences of identical length, centered on cytosines or adenines that lacked methylation evidence. Sequences with over 80% pairwise identity were removed.

The final 5mC dataset comprised 4,688 sequences, and the 6mA dataset contains 36,670 sequences, with a 1:1 positive-to-negative ratio. Each dataset was then randomly split into training (80%), validation (10%), and test sets (10%), preserving the class balance in all subsets.

#### Transcription regulation

##### Transcription factor binding site prediction

We downloaded ChIP-seq data (narrowPeak files, GRCh38) from ENCODE (version 230822), collecting 3,572 raw files. The construction of dataset was identical to DNA accessibility data. The final dataset comprised 3,572 features, with 1,428,717 sequences in the training set, 277,799 in the validation set, and 369,087 in the test set.

##### Promoter identification

We obtained promoter datasets for proximal and core promoter prediction from DNABERT-2^7^. Sequences were stratified into TATA and non-TATA groups based on the presence or absence of the TATA-box.

For proximal promoter prediction, positive sequences (300 bp; -249 bp to +50 bp relative to the transcription start site [TSS]) were derived from EPDnew. Negative sequences were sampled from non-promoter genomic regions while matching the corresponding motif constraints. The core promoter dataset was constructed similarly but used 70-bp sequences centered on the TSS (−34 to +35 bp).

The final datasets comprised three subsets: the TATA dataset (*n* = 6,130), the non-TATA dataset (*n* = 53,066) and the combined All dataset (*n* = 59,196). All datasets were randomly split into training (80%), validation (10%), and test sets (10%).

##### Enhancer identification

We obtained enhancer sequences for human and mouse from the VISTA Enhancer Browser (version 241106)^40^. Positive sequences corresponded to experimentally validated forebrain enhancers at embryonic day 11.5; negative sequences were defined as genomic regions with enhancer-like features but lacking validated activity. All sequences were partitioned into 200-bp fragments, and sequences with more than 80% pairwise identity were removed using CD-HIT.

The final human enhancer dataset comprised 12,693 sequences (3,586 positive; 9,053 negative), and the mouse enhancer dataset contains 13,378 sequences (2,639 positive; 10,739 negative). Each dataset was then randomly split into training (80%), validation, and test sets(10%), preserving the class balance in all subsets.

##### Silencer identification

Silencer data were obtained from Doni *et al.* (2020)^41^ and processed followed the DeepSilencer pipeline (https://github.com/xy-chen16/DeepSilencer). Briefly, Uncharacterized *cis*-regulatory elements (CREs) were defined by excluding genomic regions with promoter or enhancer chromatin marks, or bound by CTCF. The activity of these uncharacterized CREs was then measured via MPRA assays in K562 cell line. The 2,000 CREs with lowest MPRA activity were considered as positive samples (silencer) and the 2,000 CREs with the highest activity as negative samples. Each CRE was 200 bp in length.

Due to the limited sample size (*n* = 4,000), the dataset was randomly split into training (70%), validation (10%), and test sets (20%), preserving the class balance in all subsets.

##### CRE activity prediction

We obtained CRE activity data from Agarwal *et al.* (2025) ^42^, which employed lentiMPRA technology to quantify the transcriptional activity of promoter elements and potential enhancer elements across multiple cell lines. To construct our benchmark datasets, we selected data from two widely used cell lines: HepG2 and K562.

The final HepG2 dataset comprised 245,852 sequences, and the K562 dataset comprised 393,328 sequences; each sequence was 230 bp in length. Each dataset was randomly split into training (80%), validation (10%), and test sets (10%), preserving the class balance in all subsets.

##### Long-range interaction prediction

We obtained long-range chromatin interaction data from Li, *et. al* (2019)^43^, which integrated Ensembl TSS annotations and FANTOM5 enhancer annotations to fine-map interactions from promoter capture Hi-C and ChIA-PET data, achieving a resolution of 1 kb. Negative pairs were sampled to match the genomic distance distribution of positive interactions.

The final datasets corresponding to six distinct cell lines, with promoter sequences of 1,000 bp and enhancer sequences of 2,000 bp. The positive-to-negative ratio was approximately 1:5 in test sets and 1:20 in training and validation sets. A summary of dataset statistics is provided in Supplementary Table 2.

#### Post-transcription regulation

##### Splice site prediction

We obtained splice site prediction datasets from Scalzitti *et al*. (2021)^44^. Positive samples were defined as 400-bp sequences centered on annotated acceptor or donor splice sites. Negative samples were extracted from exon, intron, or other genomic regions containing GT/AG dinucleotides that did not function as true splice sites, matched at a 1:1 ratio.

The acceptor set comprised 22,154 sequences and the donor set comprised 21,945 sequences. Each dataset was randomly split into training (70%), validation (10%), and test sets (20%), preserving the class balance in all subsets.

##### Exon skipping rate prediction

We obtained exon skipping rate data from Cheng, *et. al.* (2021)^45^ and processed data following the methods described by Tang, *et. al* (2025)^16^. Briefly, exons from chromosomes 4, 6, 8, 10–23, X and Y were considered as training set; chromosomes 1, 7, and 9 as validation set; chromosomes 2, 3, and 5 as test set. For validation and test sets, specific filtering criteria were applied: exons were retained only if they contained missing Percent Spliced-In (PSI) values in at least 10 tissues and exhibited a PSI variance of at least 0.2 across tissues.

Each sample consisted of two 400-bp sequences, and the prediction target was the PSI profile across 56 tissues. The final dataset comprised 38,028 training samples, 1,088 validation samples, and 1,621 test samples.

##### Intron retention prediction

We obtained intron retention dataset from Daound, *et. al.*(2025)^46^. Positive samples were defined as DNA open chromatin regions associated with intron retention events, while negative samples were open regions without such events.

The final dataset comprised 75,992 sequences (600 bp), with a positive-to-negative ratio of approximately 1:2. The dataset was randomly split into training (80%), validation (10%), and test sets (10%).

##### Polyadenylation signal prediction

We obtained polyadenylation signal (PAS) data from Kalkatawi, *et. al.* (2019)^47^. Positive samples were defined as 600-bp sequences centered on annotated PAS sites. Negative sequences were extracted from chromosome 21, matched for GC content and motif composition but unrelated to polyadenylation. The dataset covered 16 signal variants, with AATAAA being the most frequent.

Two subsets were constructed: the AATAAA dataset (*n* = 22,602) and the All-variant dataset (*n* = 41,864). Each dataset was randomly split into training (60%), validation (15%), and test sets (25%), preserving the class balance in all subsets.

##### Translation initiation site prediction

The translation initiation site (TIS) dataset was also obtained from Kalkatawi, *et. al.* (2019)^47^, constructed following the identical protocol to that used for PAS datasets.

The final dataset comprised 56,488 sequences, and was randomly split into training (60%), validation (15%), and test sets (25%).

#### Gene expression prediction

##### Bulk RNA-seq

The dataset for bulk RNA-seq gene expression prediction was derived from ExPecto^48^. Briefly, each sample corresponded to a protein-coding or lncRNA gene, represented by a 2,000-bp sequence centered on its TSS. The prediction target was the log-transformed RPKM expression profiles across 53 GTEx tissues.

Genes from chromosome 8 were considered as test set (*n* = 987); chromosome 9 as validation set (*n* = 939); all other chromosomes as training set (*n* = 21,774).

#### Variant effect prediction

##### Disease-related variants

We assembled a diseased related set of single nucleotide variants (SNVs) from ClinVar^49^ (version 240307, GRCh38). To ensure data quality, we only retained SNVs located on autosomal and sex chromosomes with a review status of at least one-star. Variants classified as “Pathogenic” were used as positive samples, and those classified as “Benign” as negative samples.

Then we annotated variants using GENCODE (v44): variants located within start codons, stop codons, or coding sequence (CDS) were designated as coding variants; all others were considered as non-coding variants. The coding variant dataset comprised 122,327 variants, and the non-coding variant dataset comprised 111,202 variants.

##### Transcription-related variants

We compiled two types of transcription-related variant sets: expression quantitative trait loci (eQTLs) and MPRA-validated variants.

The eQTL datasets were derived from Enformer^14^, which characterized eQTLs across 49 distinct tissues from the GTEx project. The sample size varies by tissue, with variant counts ranging from 108 to 5,480.

For MPRA dataset, we randomly sampled 210,000 variants tested in K562 and HepG2 cell line from REVA^50^, including 10,000 transcription-modulating variants and 200,000 non-modulating variants.

### Benchmark models

#### Genomic language models

We conducted a systematic evaluation of 11 genomic language models. These models were selected for their diversity in pretraining data, tokenization strategies, architectures, and pretraining objective. A detailed summary for each model is provided in Supplementary Table 3.

#### Baseline models

##### Non-gLM baselines

We primarily selected non-gLM baseline models from well-established published models that were not trained with genomic language models. For specific applications, we retrained convolutional neural network-based models following established practices as baseline models (Supplementary Table 4).

For DNA accessibility prediction, histone modification prediction, and transcription factor binding site prediction, we constructed convolutional neural network baseline models following the Basset^38^ architecture (Supplementary Fig. 2). Models were trained with a batch size of 128, using binary cross entropy (BCE) as the loss function, the Adam optimizer (with default parameters). Early stopping was applied based on validation loss. These models were implemented in Python using the TensorFlow deep learning framework (https://www.tensorflow.org/).

For DNA methylation prediction, we constructed similar convolutional neural network baseline models with simplified architecture (Supplementary Fig. 3) due to the relatively small size of the training datasets. The training strategy and hyperparameters were identical to those described previously. These models were implemented in Python using the PyTorch deep learning framework.

##### Random baseline

To isolate the contribution of pretraining, we constructed BERT-based random models: RandomWeight. RandomWeigh model contained 12 encoder layers, 1,024 hidden dimensions, and 155M parameters. It was initialized randomly with a maximum input length of 4,096 (single-nucleotide tokenization) and evaluated without any training (Supplementary Fig. 4).

### Benchmarking methods

To evaluate the information captured during pretraining, we assessed each gLM using either adapter fine-tuning or zero-shot strategy (Supplementary Fig. 5). For sequence embedding, we used the final layer output for all models except Evo2, for which we followed the original authors’ recommendation to use the 29th layer’s output.

#### Downstream model architecture

A unified CNN-based architecture was used for most applications (Supplementary Fig. 6). Outputs were activated by a sigmoid function for classification and a linear function for regression.

For DNA accessibility, histone modification, and transcription factor binding site prediction, which involving a large number of output features, we adopted the same model architecture as the corresponding non-gLM baseline (Supplementary Fig. 2) to ensure effective feature extraction. All training settings remained consistent with the baseline.

For downstream applications requiring dual-sequence input, such as long-range interaction prediction and exon skipping rate prediction, we designed customized architecture based on the general downstream model (Supplementary Fig. 7). In particular, the model for exon skipping rate prediction closely followed the architecture described in the corresponding original publication^45^ (Supplementary Fig. 8).

#### Model training

Downstream models were trained with a batch size of 128 using Adam optimizer (default parameters) and early stopping strategy. The loss function was selected as BCE for classification task and mean squared error (MSE) for regression task. All models were implemented in Python using the PyTorch deep learning framework.

#### Variant effect prediction

For zero-shot variant effect prediction, 512-bp sequences centered on each variant were extracted for both reference and alternative alleles. Sequence embeddings were generated and averaged across all nucleotide positions. A variant impact score was then calculated as the Euclidean distance between the reference and alternative sequence embeddings (Supplementary Fig. 9).

#### Evaluating metrics

Model performance was evaluated using task-appropriate metrics. For binary classification tasks, the area under the receiver operating characteristic curve (AUROC) was used. For regression tasks, performance was assessed using the Spearman correlation coefficient.

### Human (model H) and multi-species (model M) gLM training

We downloaded reference genomes for 7 species (Supplementary Table 5) from GenBank. Each genome was segmented into non-overlapping 4096-bp bins, and bins containing >5% ambiguous (N) bases were excluded. We trained two models: a human gLM using only the human reference genome, and a multi-species gLM using all 7 genomes. Both models shared an identical architecture with RandomWeight and were trained using masked language modeling with a 15% masking ratio (Supplementary Fig. 10). Training was performed using the AdamW optimizer (β1 = 0.9, β2 = 0.98, ε = 1 × 10⁻⁶) with early stopping. The learning rate followed a linear schedule, warming up from 1 × 10⁻⁶ to 1 × 10⁻⁴ over 80,000 steps and then decaying linearly to a minimum of 1 × 10⁻⁶ by 2,000,000 steps. Model **H** was trained with a batch size of 16, whereas the model **M** used a batch size of 64. All models were implemented in Python using the PyTorch deep learning framework and trained on NVIDIA Tesla A100 GPUs.

### Assessing alignment between pretraining and downstream genomic applications

To quantify the contribution of pretraining, we assessed the alignment by fitting a linear regression between the pretraining loss (across training epochs) and downstream task performance, using checkpoints from model **M**. The degree of alignment was represented by the slope of this regression: a strong negative slope indicates that lower pretraining loss corresponds to substantially improved downstream performance, reflecting a well-aligned task. In contrast, a positive or near-zero slope suggests that pretraining optimization does not benefit—or even adversely affects—downstream performance, indicating misalignment between the pretraining objective and the target application.

### An information-theory perspective on the learning mechanism of MLM objective in gLMs

Consider a gLM trained on genomic sequences from *N* species, where the empirical data distribution *P*_*data*_ approximates an underlying evolutionary joint distribution *P*(*x*^(1)^, … *x*^(*N*)^) that encodes both phylogenetic relationships and sequence-level functional constraints. Pretraining proceeds via MLM, with the objective of minimizing the following loss function:

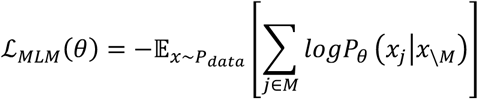

where *M* denotes the set of masked position indices and *x*_\*M*_ represents the unmasked context sequence.

From the perspective of information theory, the MLM loss corresponds to the expected cross-entropy between the model’s predictive distribution *P*_*θ*_ and the true conditional distribution *P*_*data*_. For any given context *x*_\*M*_, let *X*_*M*_ denote the masked tokens viewed as random variable and *X*_\*M*_ the unmasked context:

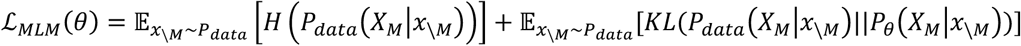

The first term represents the conditional entropy *H*_*data*_(*X*_*M*_|*X*_\*M*_), an intrinsic dataset property that remains constant, and the second term represents the expected Kullback-Leibler (KL) divergence, quantifying the discrepancy between the model-learned and true conditional distributions.

Since

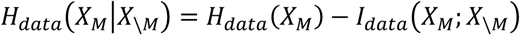

is an invariant, minimizing the MLM loss is equivalent to minimizing the KL divergence. Consequently, the mutual information captured by the model, *I*_*θ*_(*X*_*M*_; *X*_\*M*_), approaches the theoretical upper bound imposed by the data, *I*_*data*_(*X*_*M*_; *X*_\*M*_):

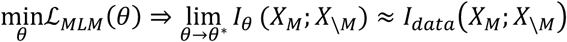

Thus, the optimization process can be viewed as maximizing the representable mutual information *I*_*θ*_(*X*_*M*_; *X*_\*M*_) bounded by the data’s intrinsic dependencies.

In the context of multiple-species genomic sequences, the mutual information *I*_*θ*_ can be further decomposed as:

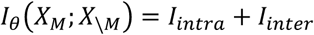

where *I*_*intra*_ denotes intra-species mutual information, capturing statistical dependencies among genomic region within individual species, and *I*_*inter*_ represents inter-species mutual information, encoding statistical dependencies between homologous sequences across different species and reflecting evolutionary constraints.

### Masked token prediction

To assess whether gLMs can capture varying levels of mutual information across human genomic regions, we randomly sampled 1,000 exon sequences from GENCODE (v44) and 1,000 Alu repeat sequences from RepeatMasker (v4.0.5)^51^. For each nucleotide position, we iteratively masked every nucleotide position and measured the predicted probability assigned to the original nucleotide. Higher probabilities indicate stronger capture of position-specific statistical dependencies.

## Data Availability

A complete list of the source datasets used is provided in Supplementary Table 6.

## Code Availability

A comprehensive list of the models and pre-trained weights evaluated in this study is provided in Supplementary Table 3. All codes for training and evaluating models are available at https://github.com/gao-lab/LingoDNABench.

## Author Contributions

G.G. and Y. W. conceived the study and supervised the research. Y.-X. L. and Y.W. designed and implemented the computational framework and proposed the theoretical analysis with guidance from G.G.. J.-C. W. participated in code testing and verification. Y.-X. L. and W.-Y. P. conducted benchmarks. Y.-X. L., Y.W., Z.-Y. C. and G.G. wrote the manuscript.

## Acknowledgments

This work was supported by funds from the National Science and Technology Major Project (grant no. 2022ZD0115004), the Changping Laboratory, the State Key Laboratory of Gene Function and Modulation Research and the Beijing Advanced Innovation Center for Genomics at Peking University. Part of the analysis was carried out on the Computing Platform of the Center for Life Sciences of Peking University and supported by the High-performance Computing Platform of Peking University and Changping Laboratory.

## Supplementary Figures

**Supplementary Fig. 1.**
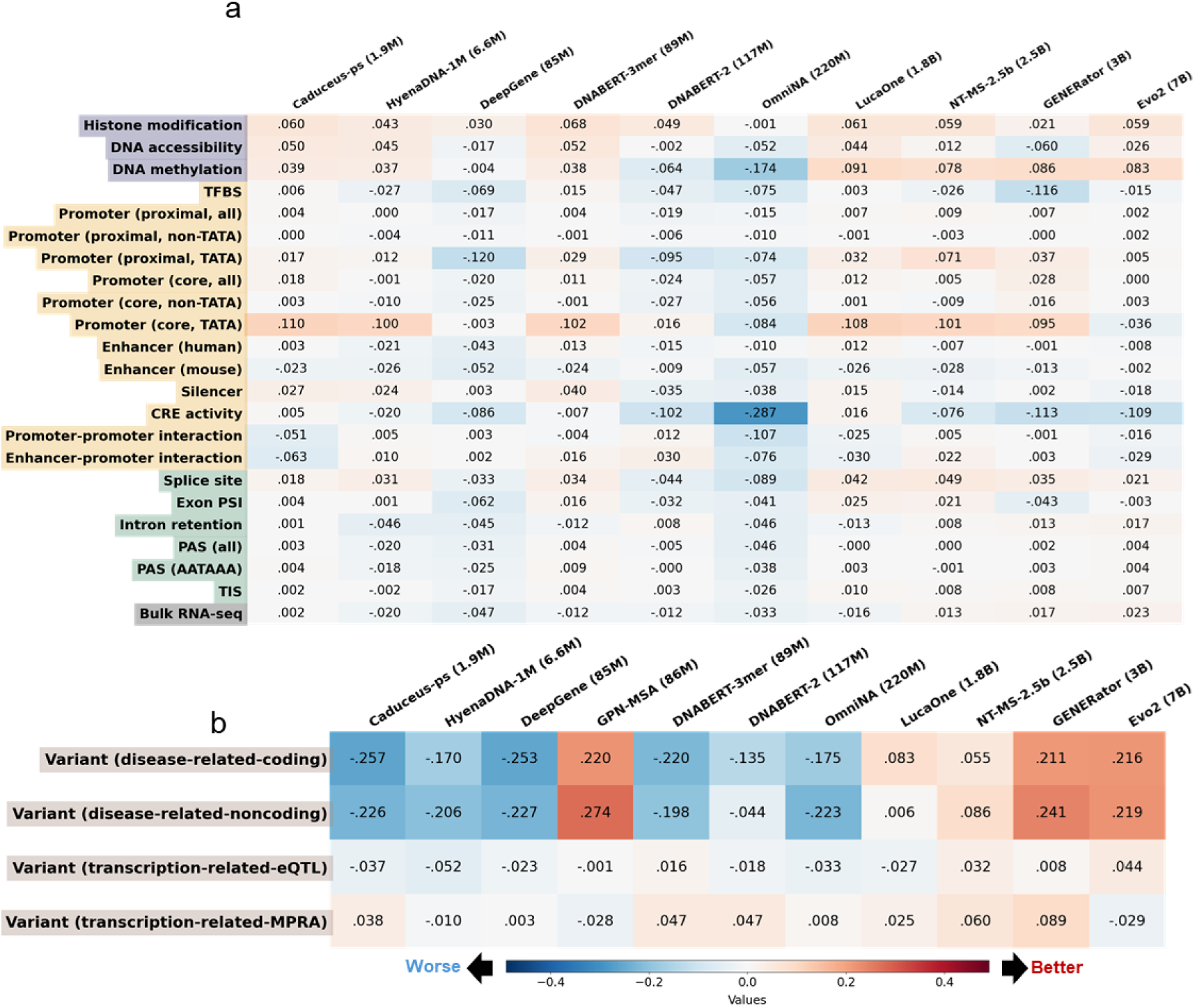
Evaluation of genomic language models on LingoDNABench. **a.** Performance gap between published gLMs and locally pretrained multi-species gLM in regulatory-oriented applications (*metric*_*gLM*_ − *metric* _*locally pretrained multi*−*species gLM*_). Red: gLMs superior; blue: locally pretrained multi-species gLM superior, same as in (b). **b.** Performance gap between published gLMs and locally pretrained multi-species gLM in variant effect prediction (*metric*_*gLM*_ − *metric* _*locally pretrained multi*−*species gLM*_).

**Supplementary Fig. 2.**
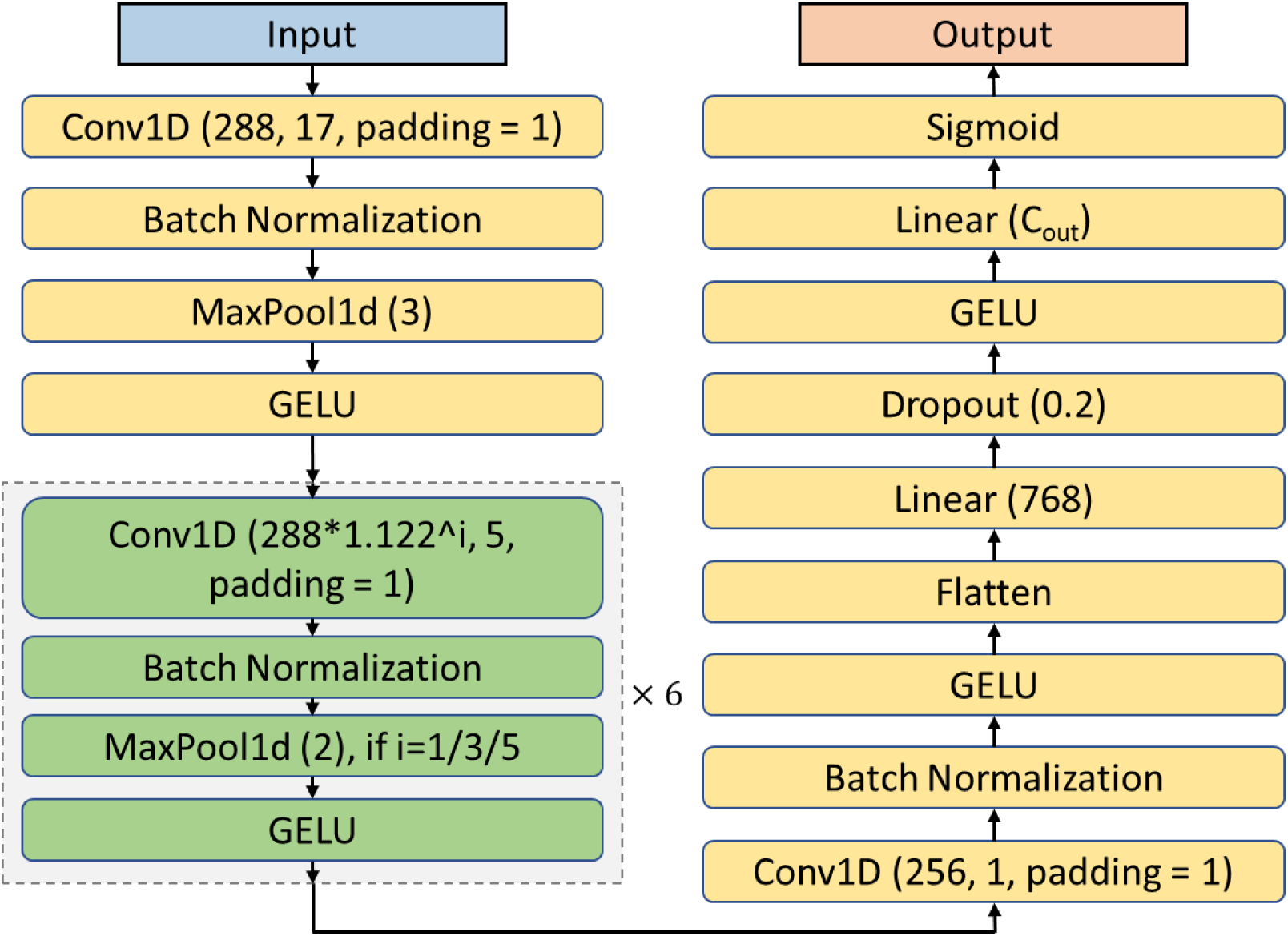
Baseline model architecture for DNA accessibility, histone modification, and transcription factor binding sites prediction. The output channel size (C_out_) is 1,770 for DNA accessibility, 3,070 for histone modification, and 3,572 for transcription factor binding sites prediction.

**Supplementary Fig. 3.**
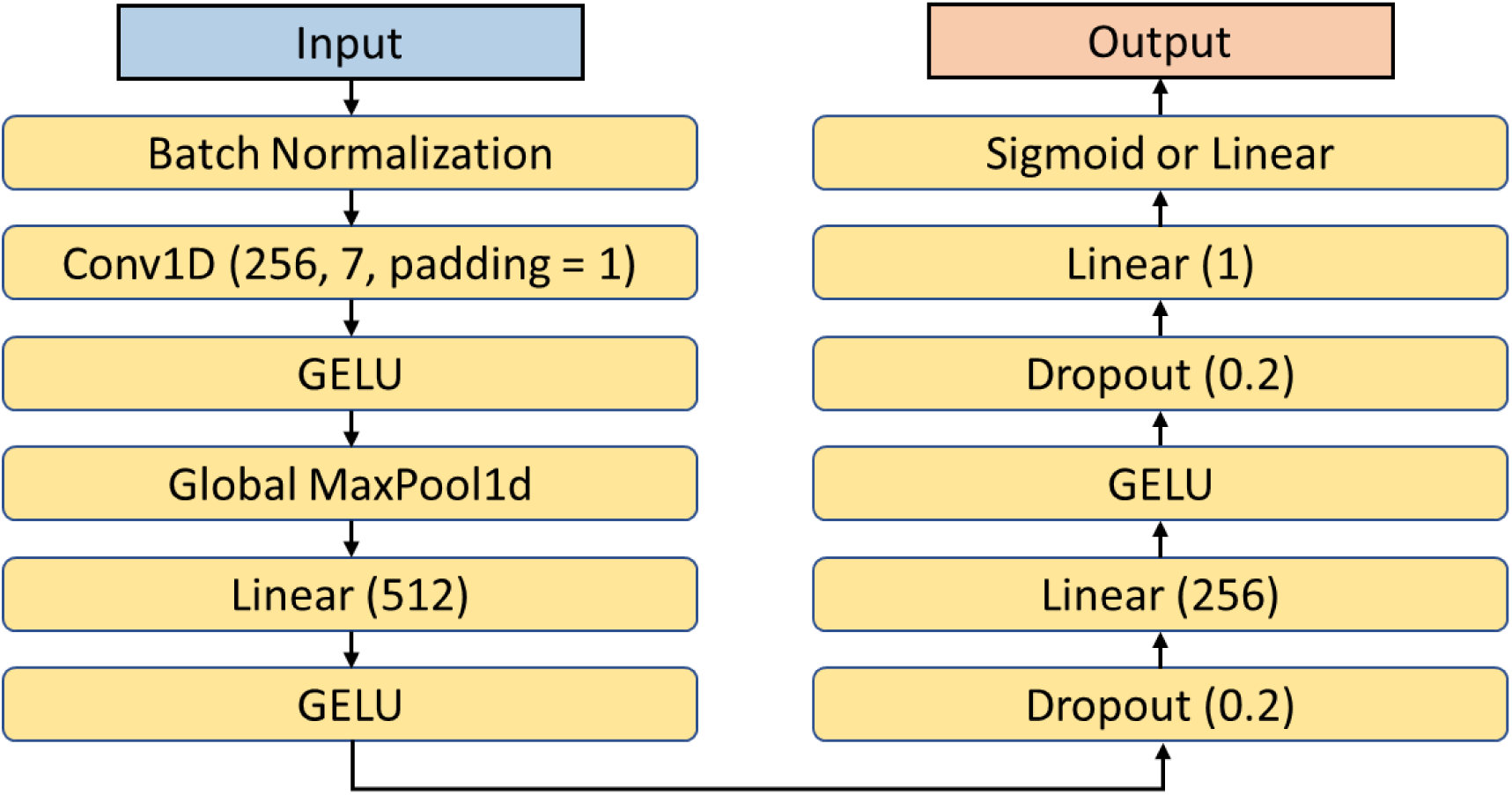
Baseline model architecture for DNA methylation and RNA Pol II elongation potential prediction. For DNA methylation prediction (binary classification), the final layer employs a sigmoid activation; for the RNA Pol II elongation potential prediction (regression), a linear activation is used. C_out_ represents the number of output features configured for each task.

**Supplementary Fig. 4.**
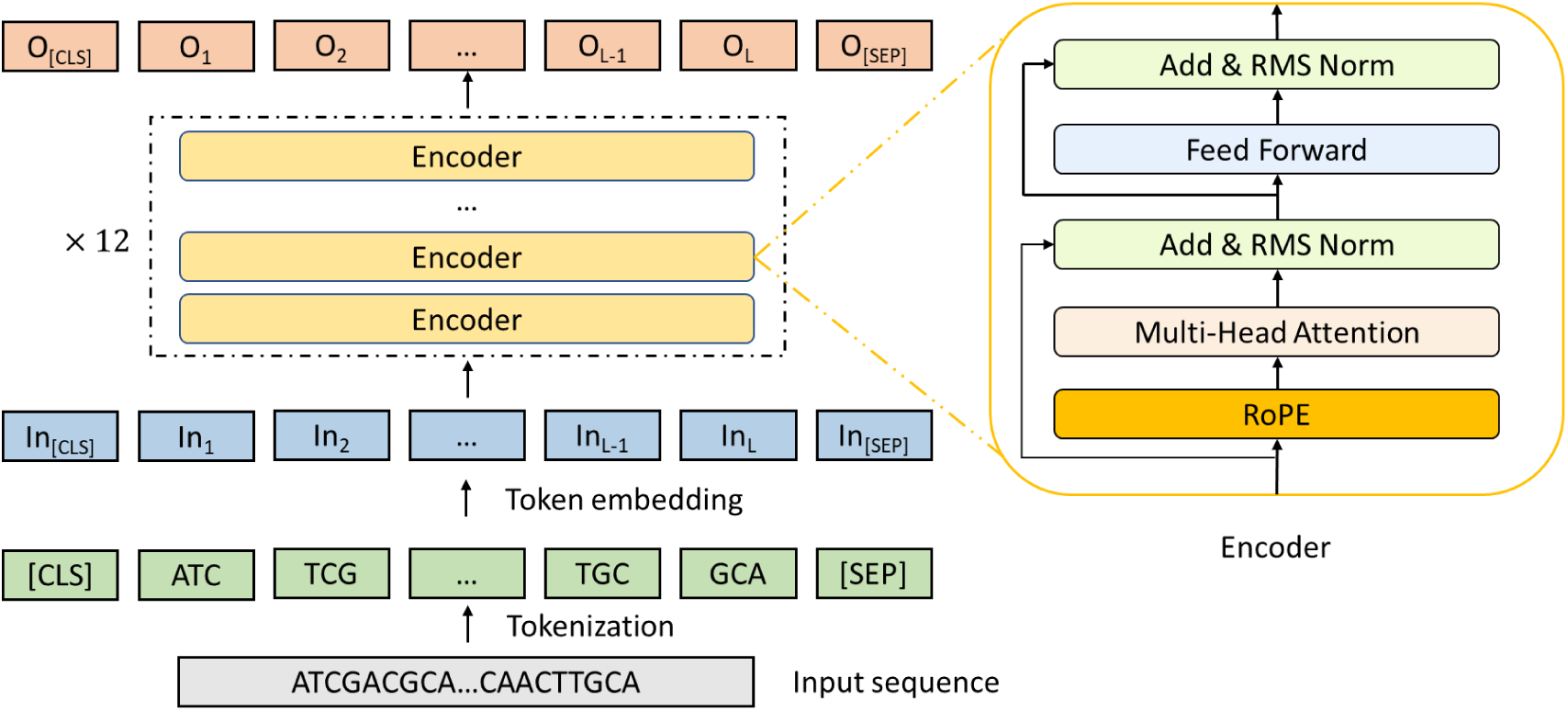
Architecture of the RandomWeight baseline model. The model is based on BERT, consisting of 12 encoder layers with a hidden dimension of 1,024 (∼155M parameters), incorporates rotary positional embedding (RoPE), and uses single-nucleotide tokenization.

**Supplementary Fig. 5.**
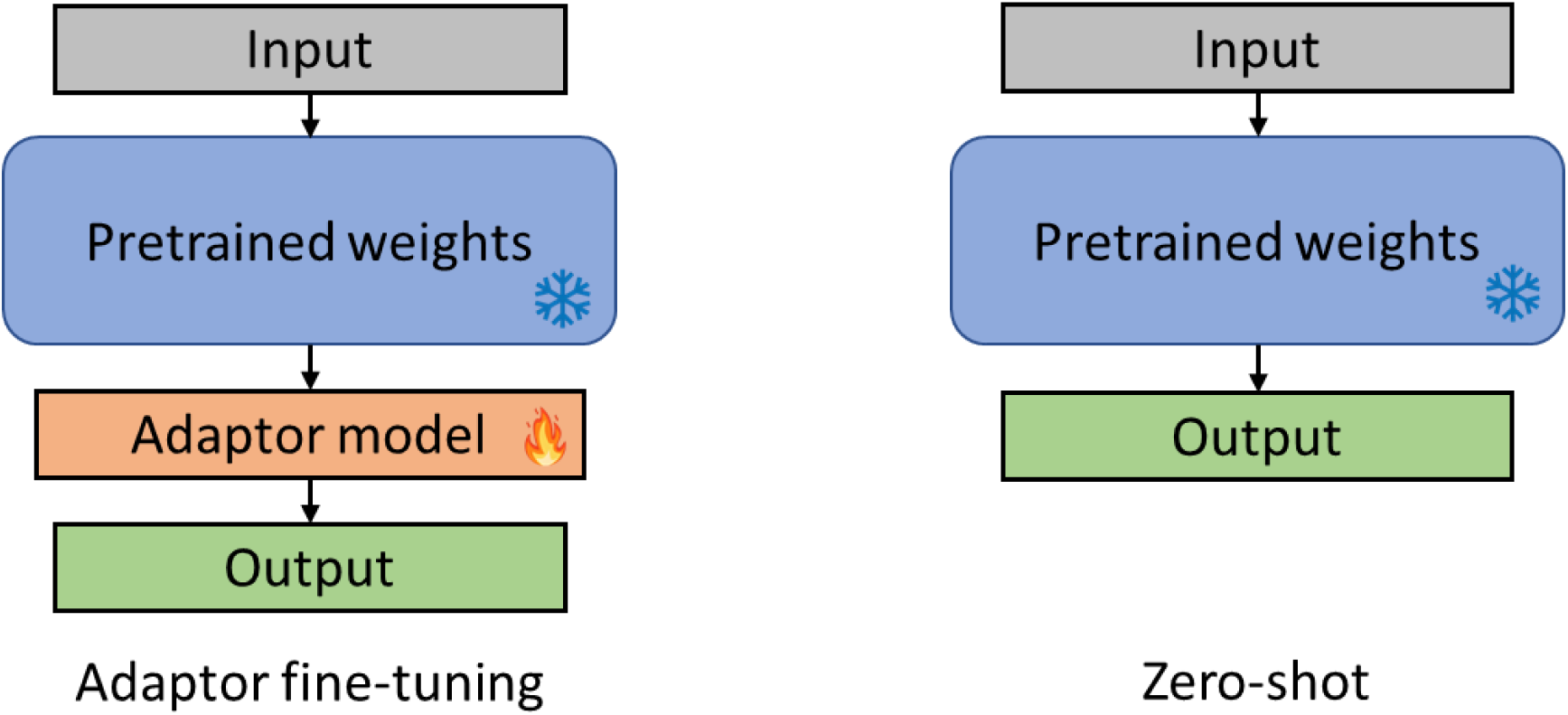
Schematic for applying pretrained genomic language models to downstream applications. The illustration of two strategies for downstream adaptation. The flame icon denotes model components with trainable parameters, whereas a snowflake icon denotes components with frozen parameters.

**Supplementary Fig. 6.**
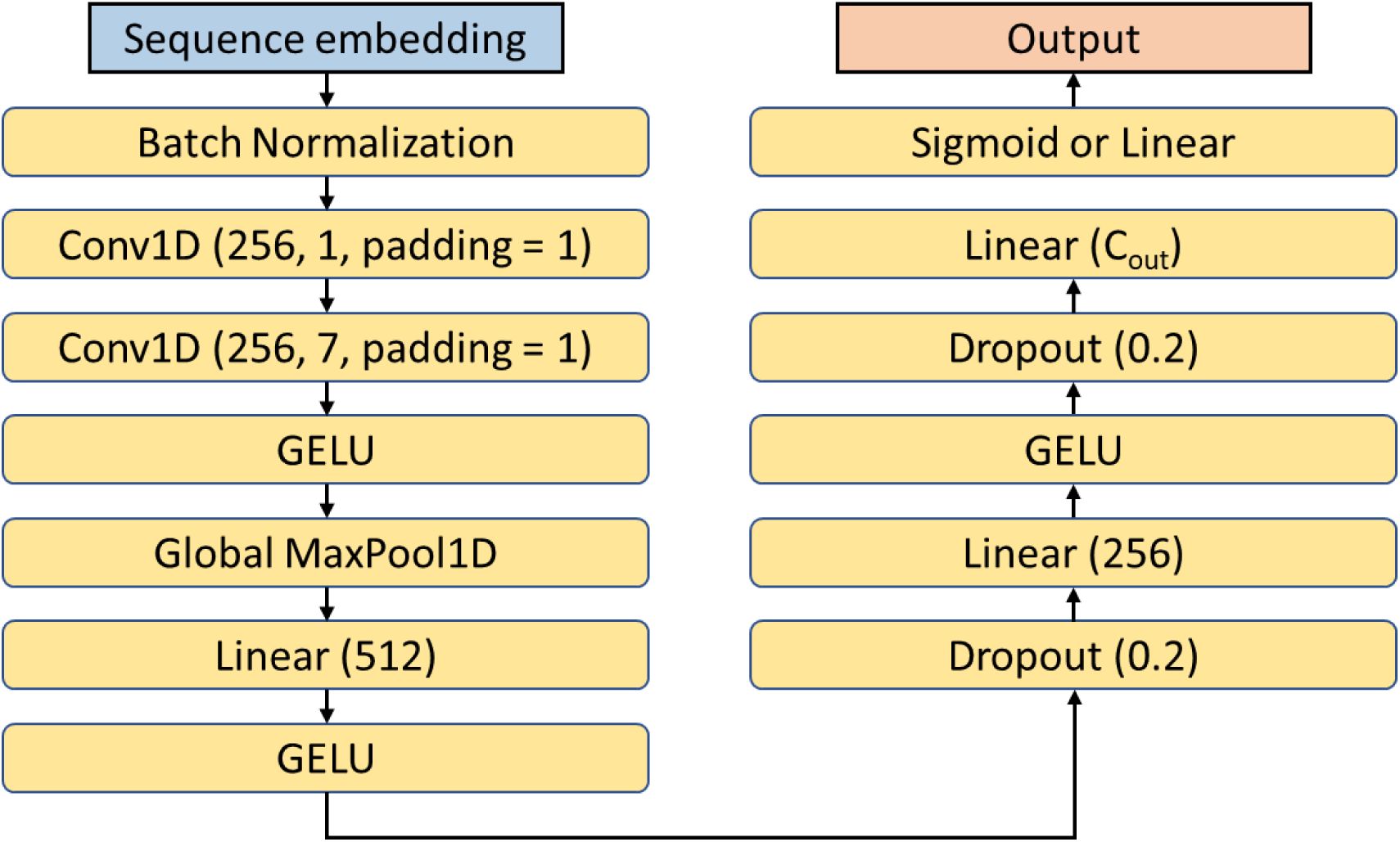
Default downstream model architecture. The CNN-based downstream model architecture was used for most applications. The output layer employs a sigmoid activation function for classification tasks and a linear activation for regression tasks.

**Supplementary Fig. 7.**
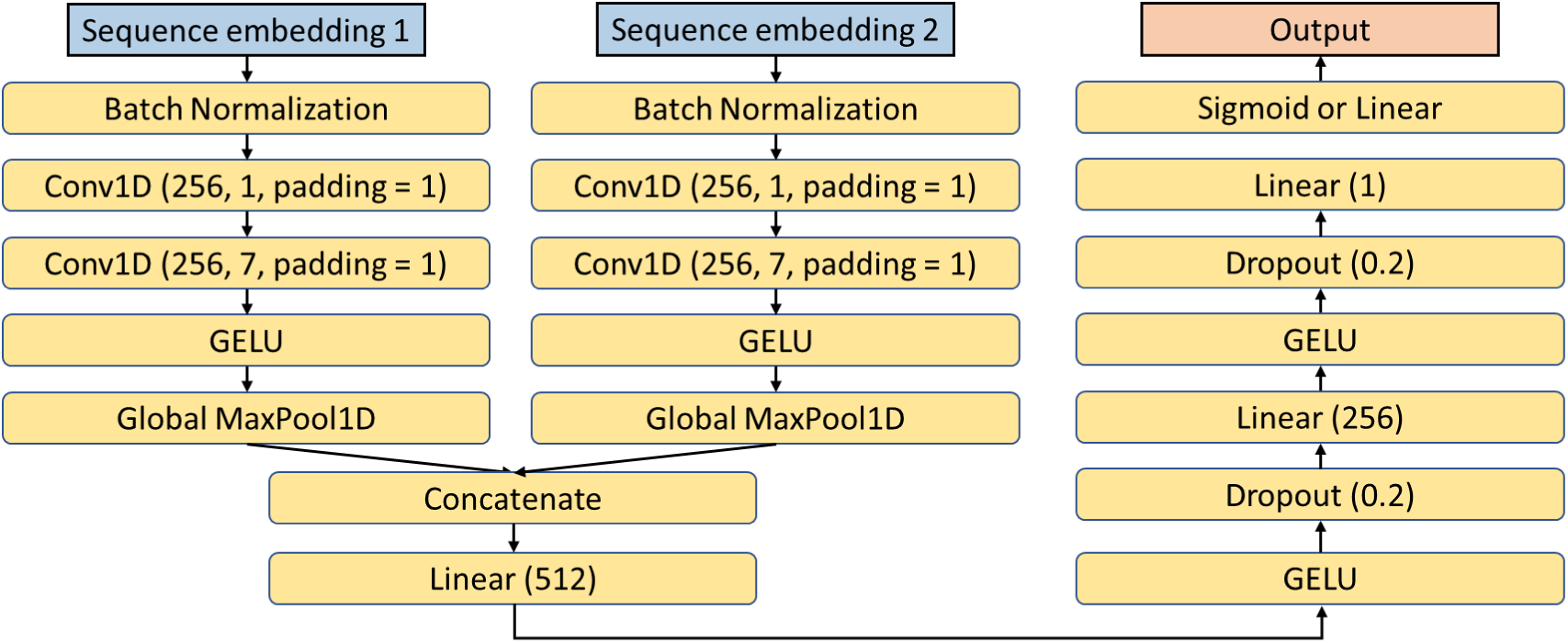
Downstream model architecture for long-range interaction prediction. The model takes two sequences as input, and the output layer uses a sigmoid activation function for binary classification.

**Supplementary Fig. 8.**
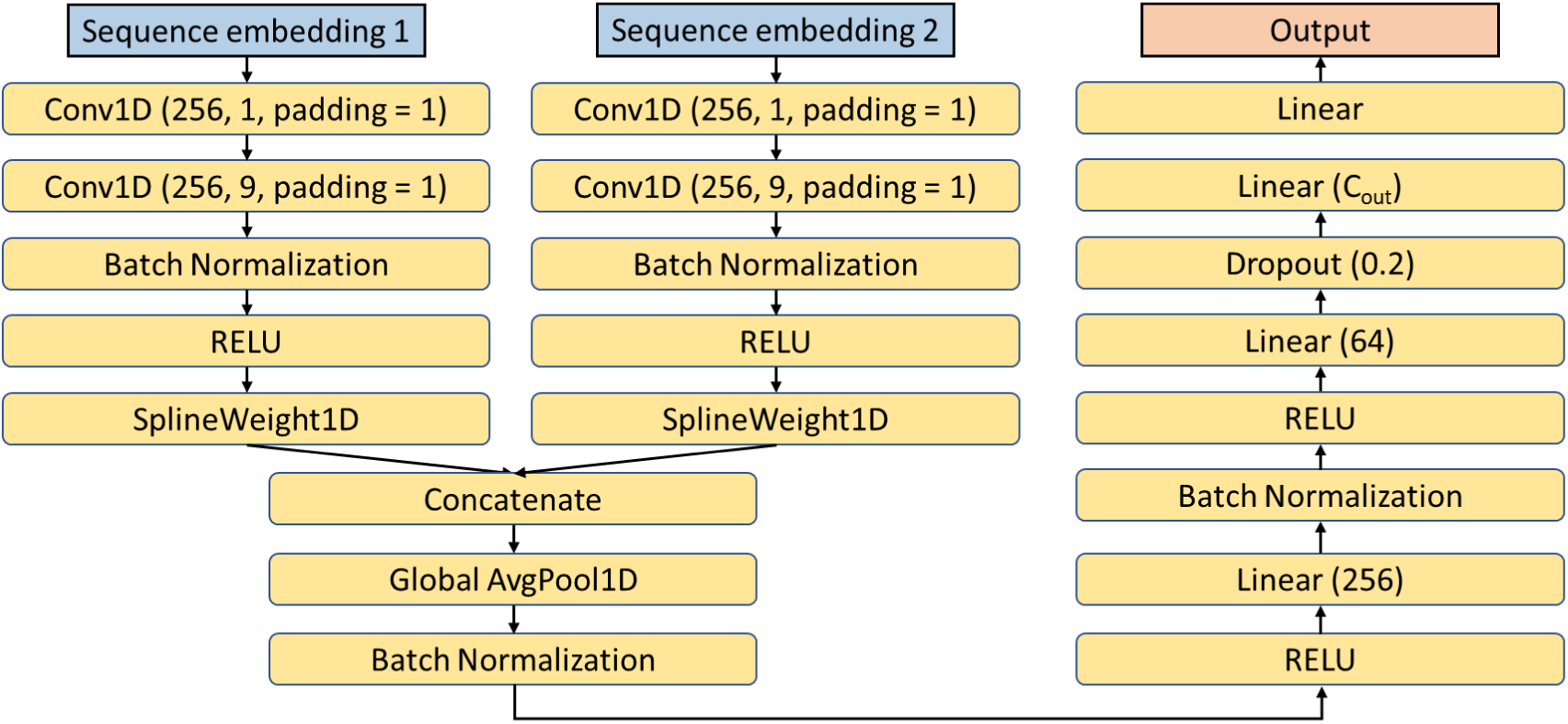
Downstream model architecture for exon skipping rate prediction. The model incorporates a custom SplineWeight1D layer, implemented with reference to the MMSplice framework (https://github.com/gagneurlab/mmsplice_mtsplice), which is specialized for modeling splicing dynamics.

**Supplementary Fig. 9.**
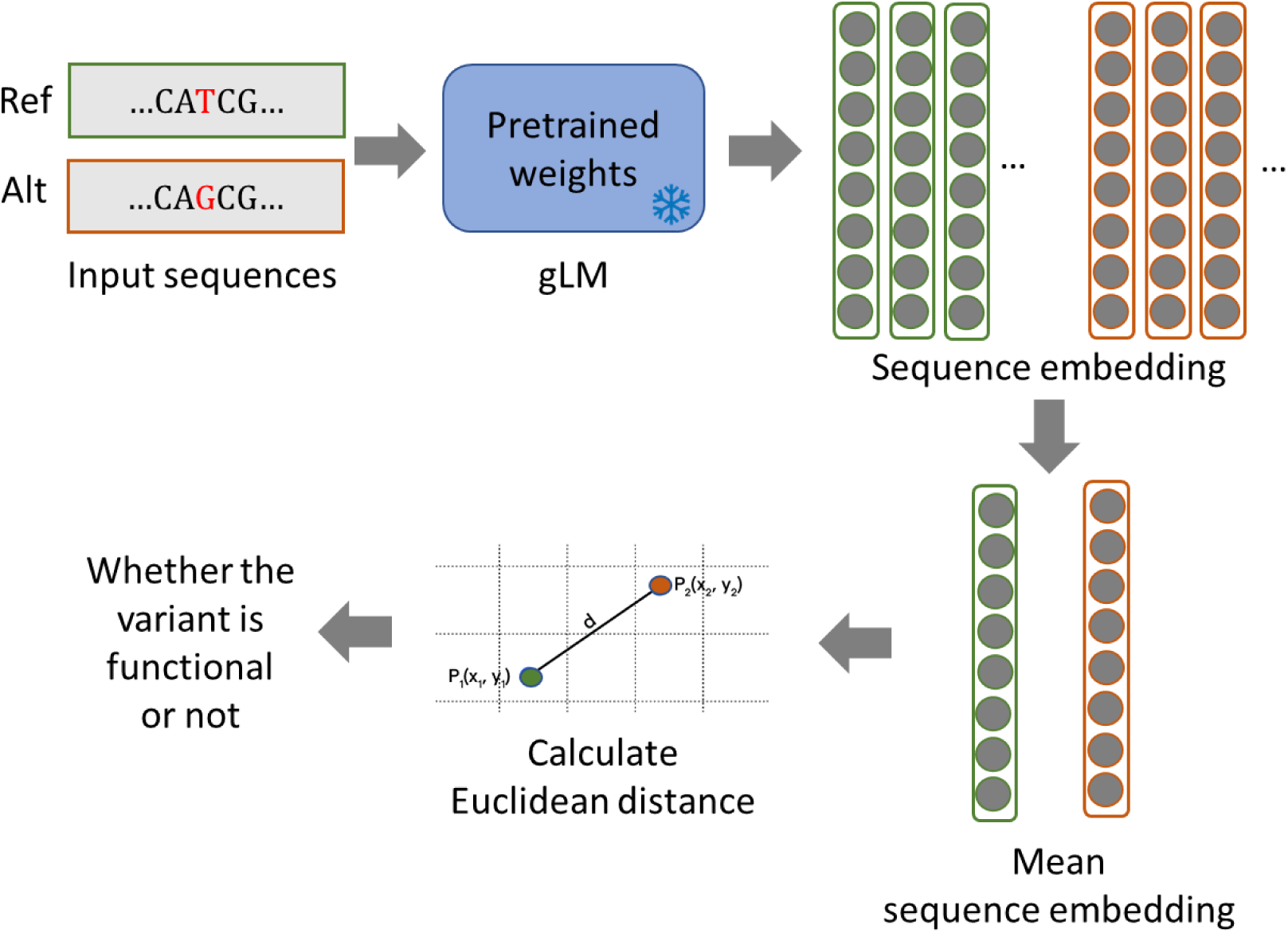
Zero-shot variant effect prediction via embedding divergence. We evaluated variant functional impact (e.g., pathogenicity or altered gene transcription) by measuring the Euclidean distance between reference and alternative allele sequence embeddings in a pretrained gLM’s latent space. The approach was based on the premise that functionally neutral variants induce minimal embedding shifts, while impactful variants (e.g., pathogenic or regulatory) cause significant divergence.

**Supplementary Fig. 10.**
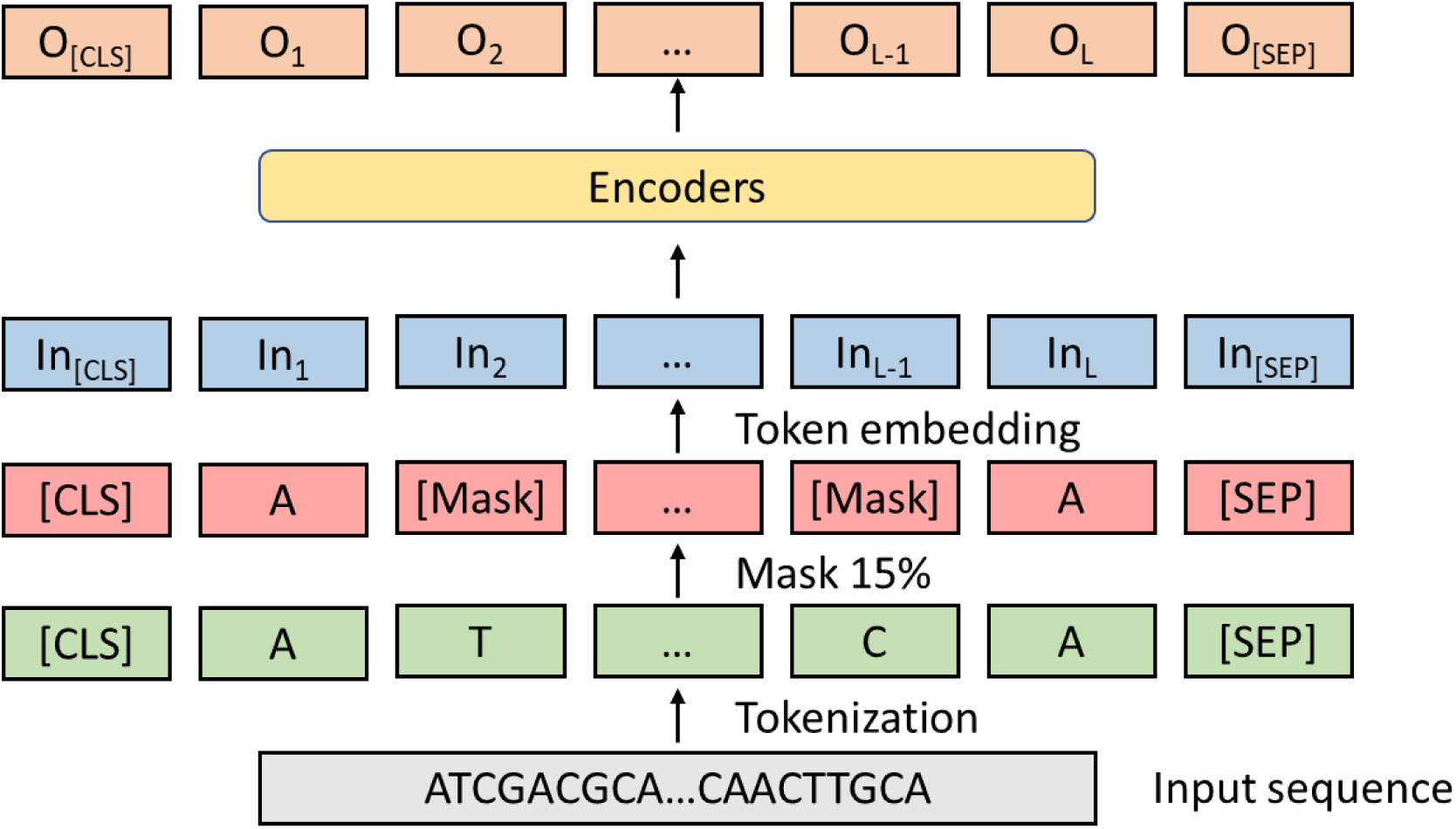
Pretraining scheme for the model H and model M. The model was pretrained using masked language modeling on 4,096-nt sequences with single-nucleotide tokenization. For each input sequence, 15% tokens are randomly masked: 80% replaced with a [MASK] token, 10% replaced with a random token, and 10% retained unchanged. Training was accelerated using FlashAttention-2.

## Supplementary Tables

**Supplementary Table 1.**
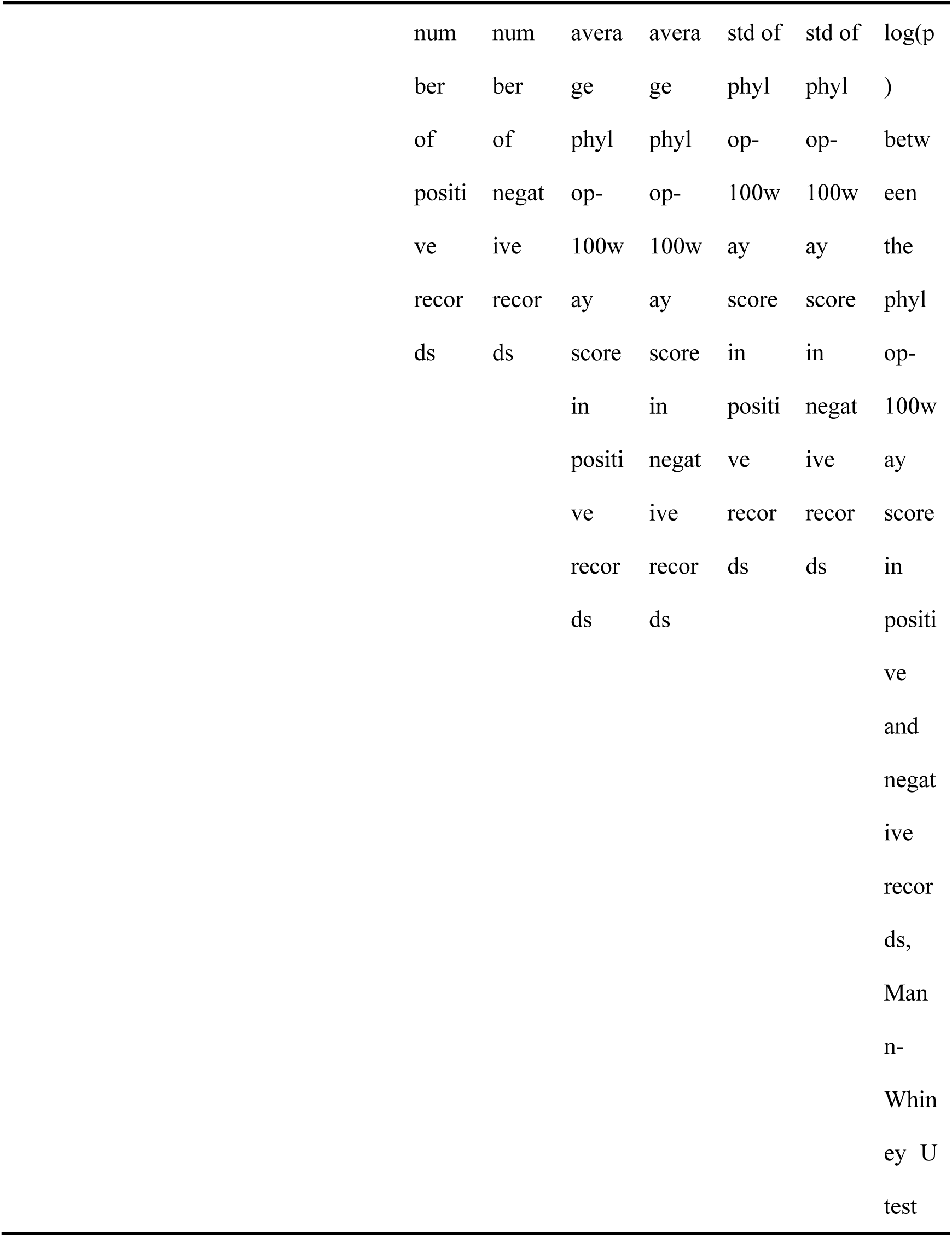
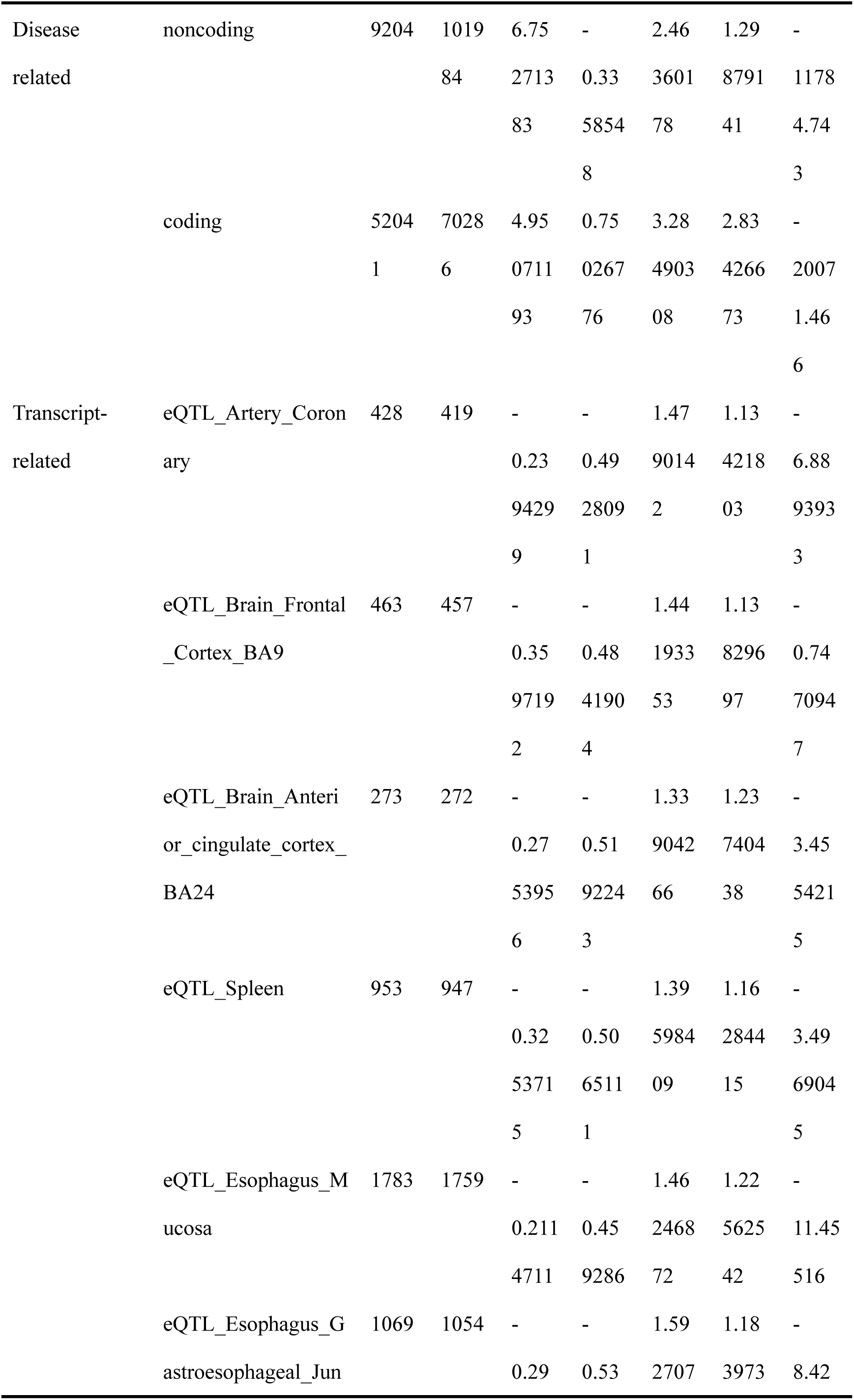
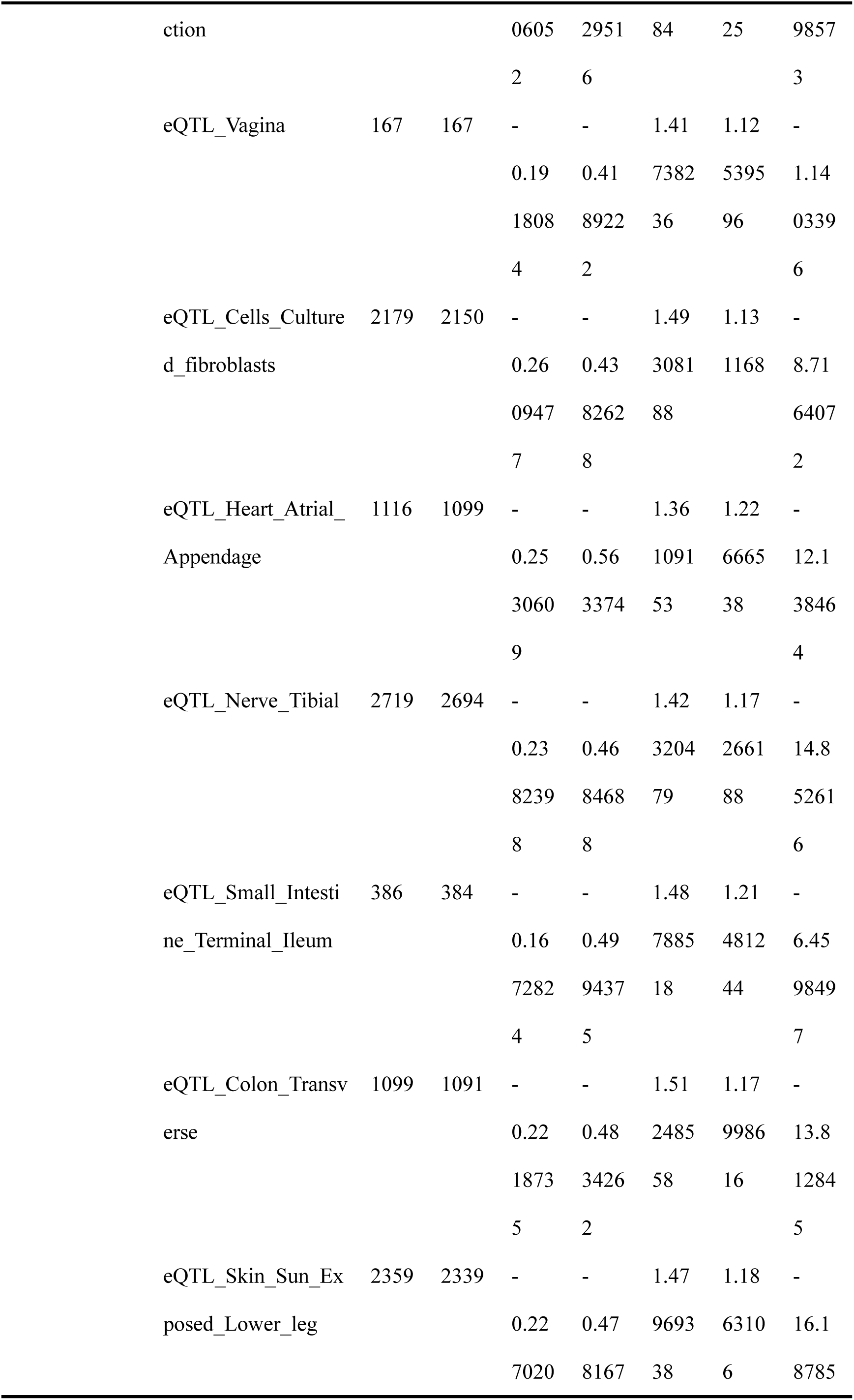
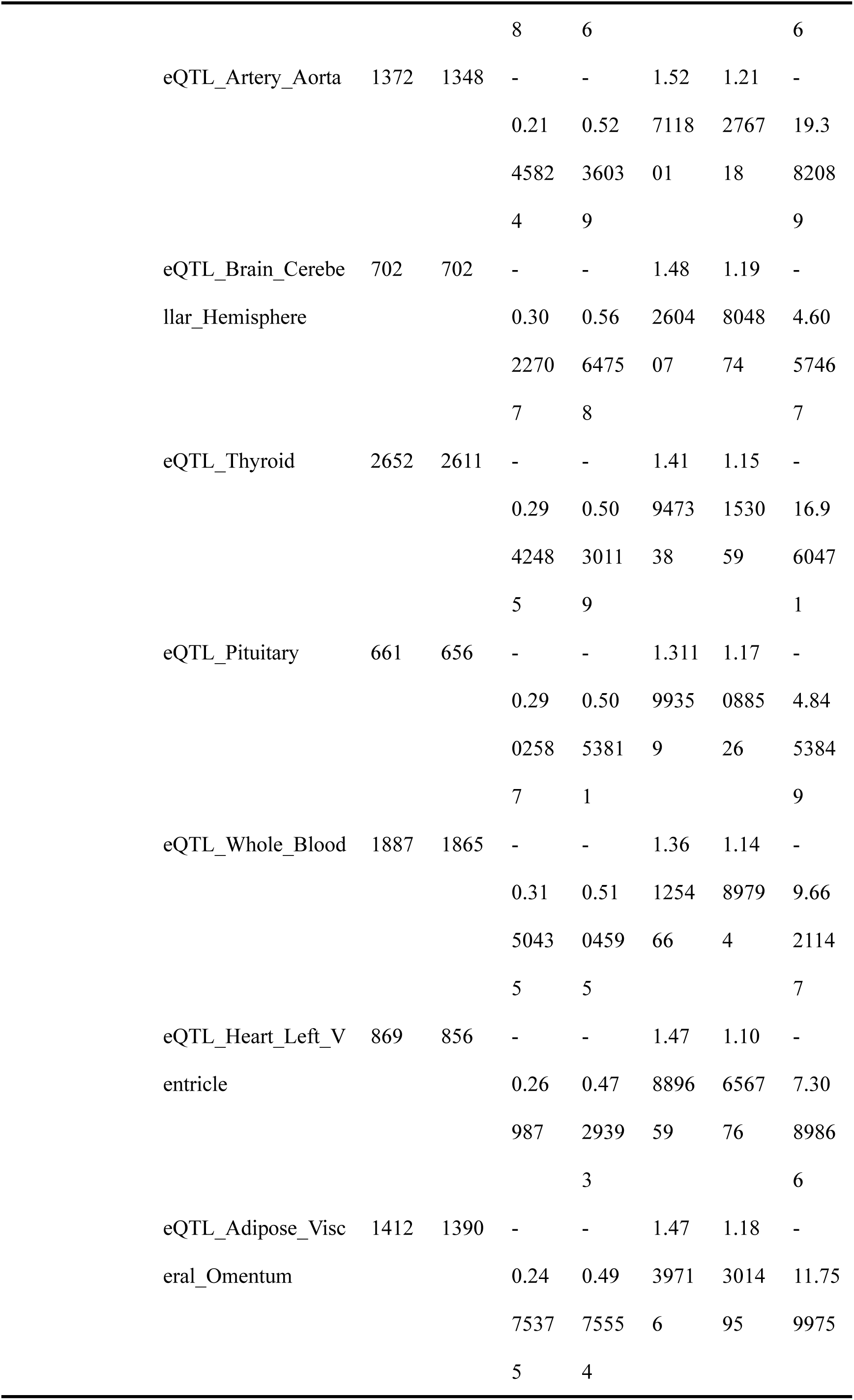
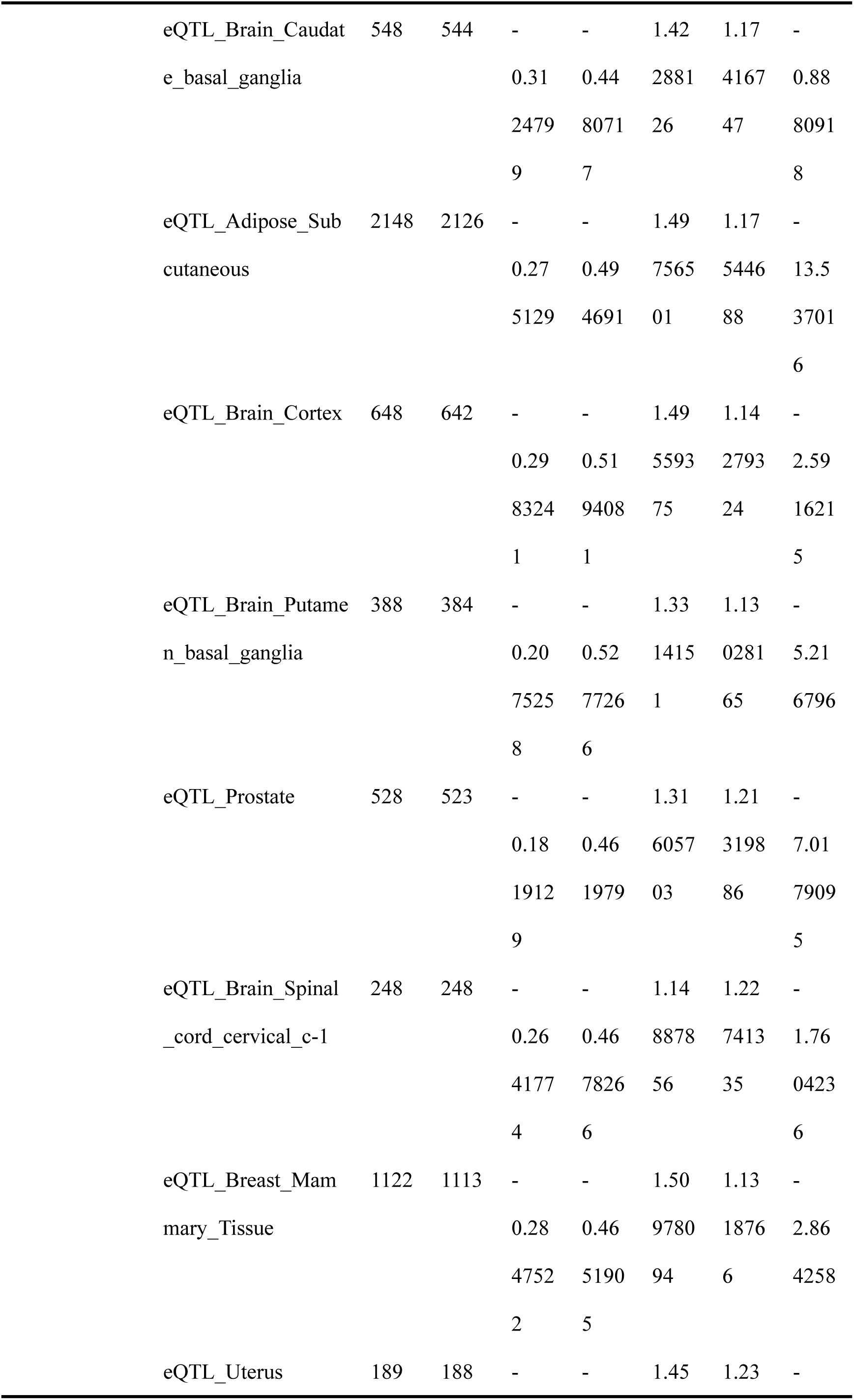
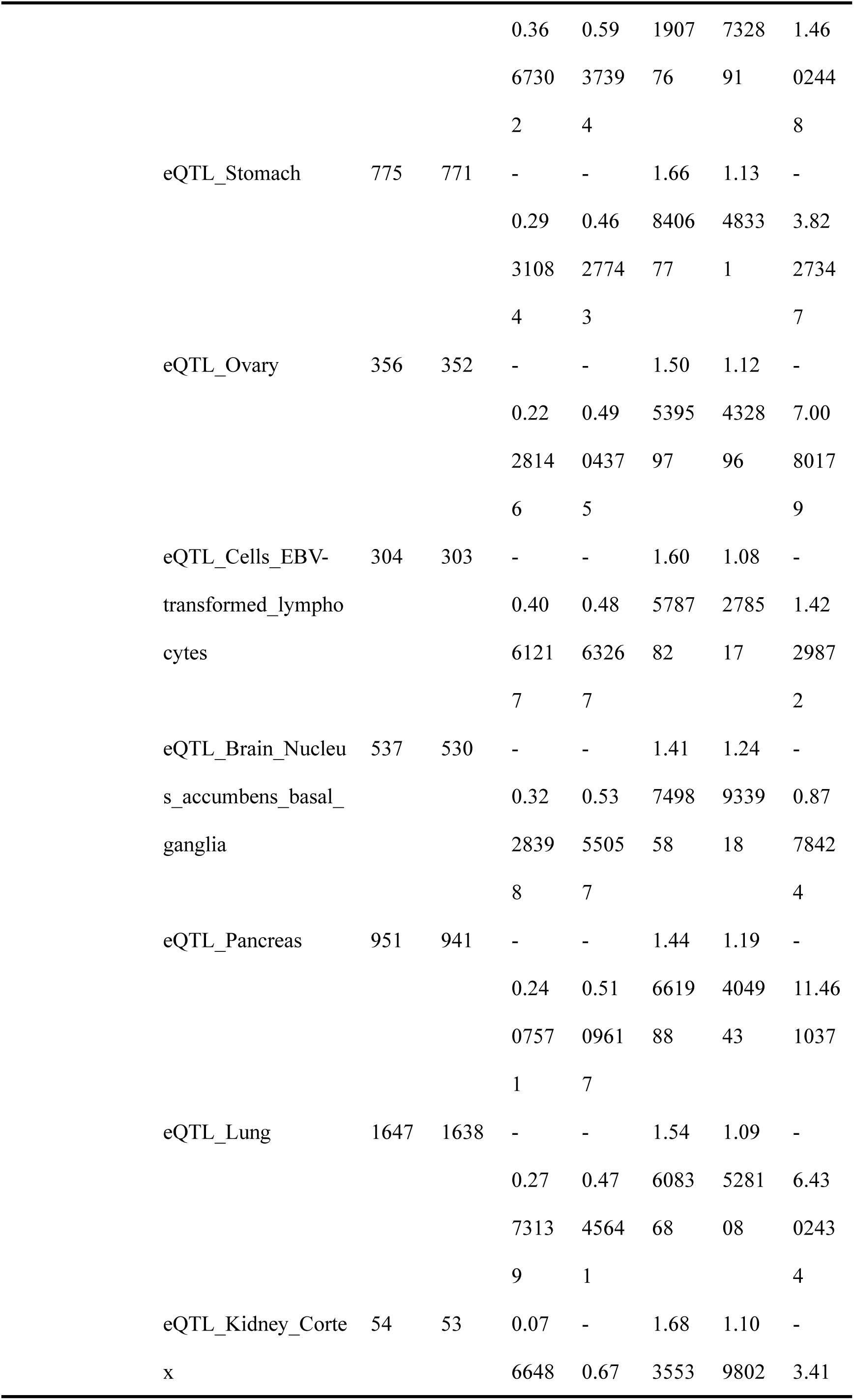
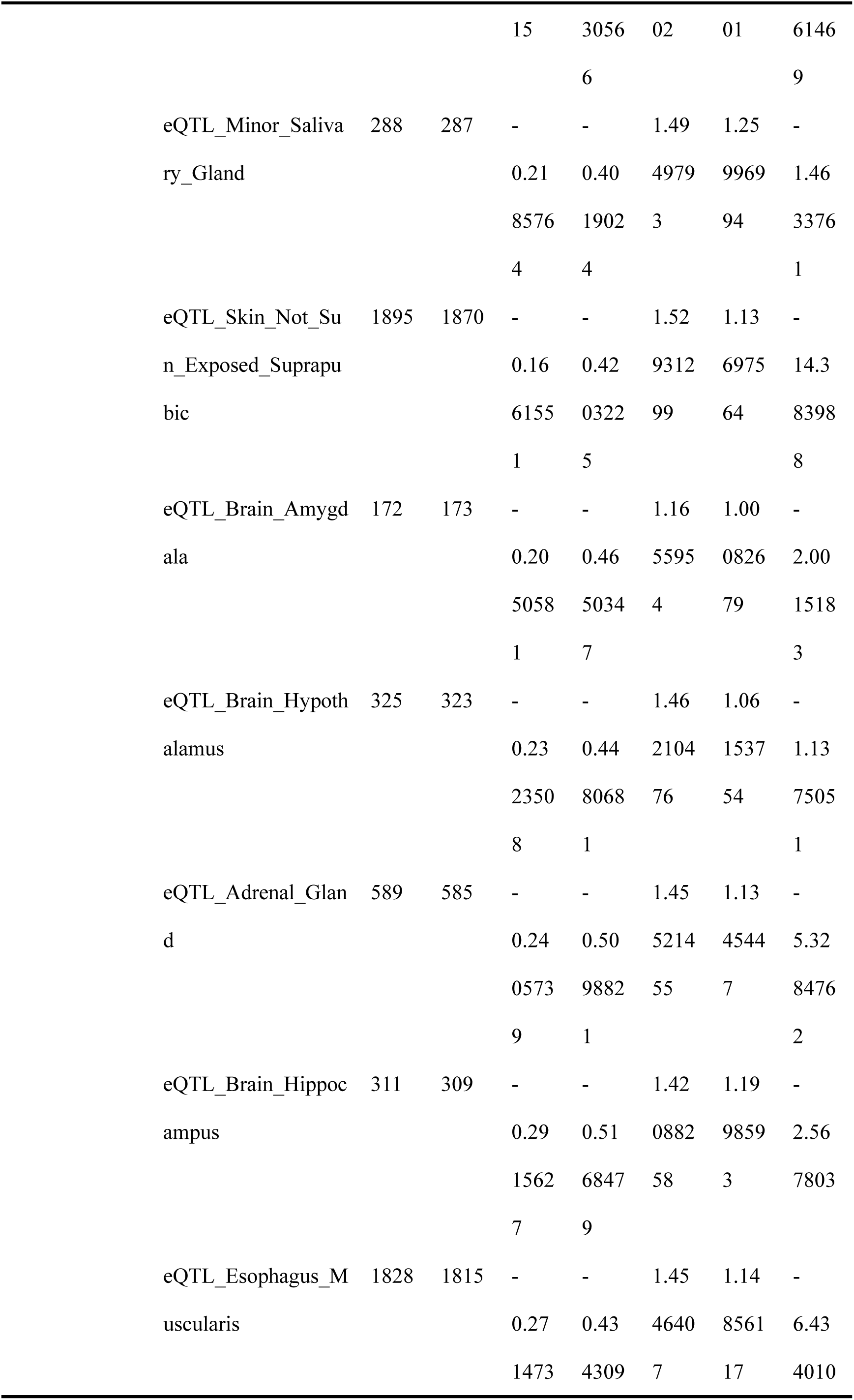
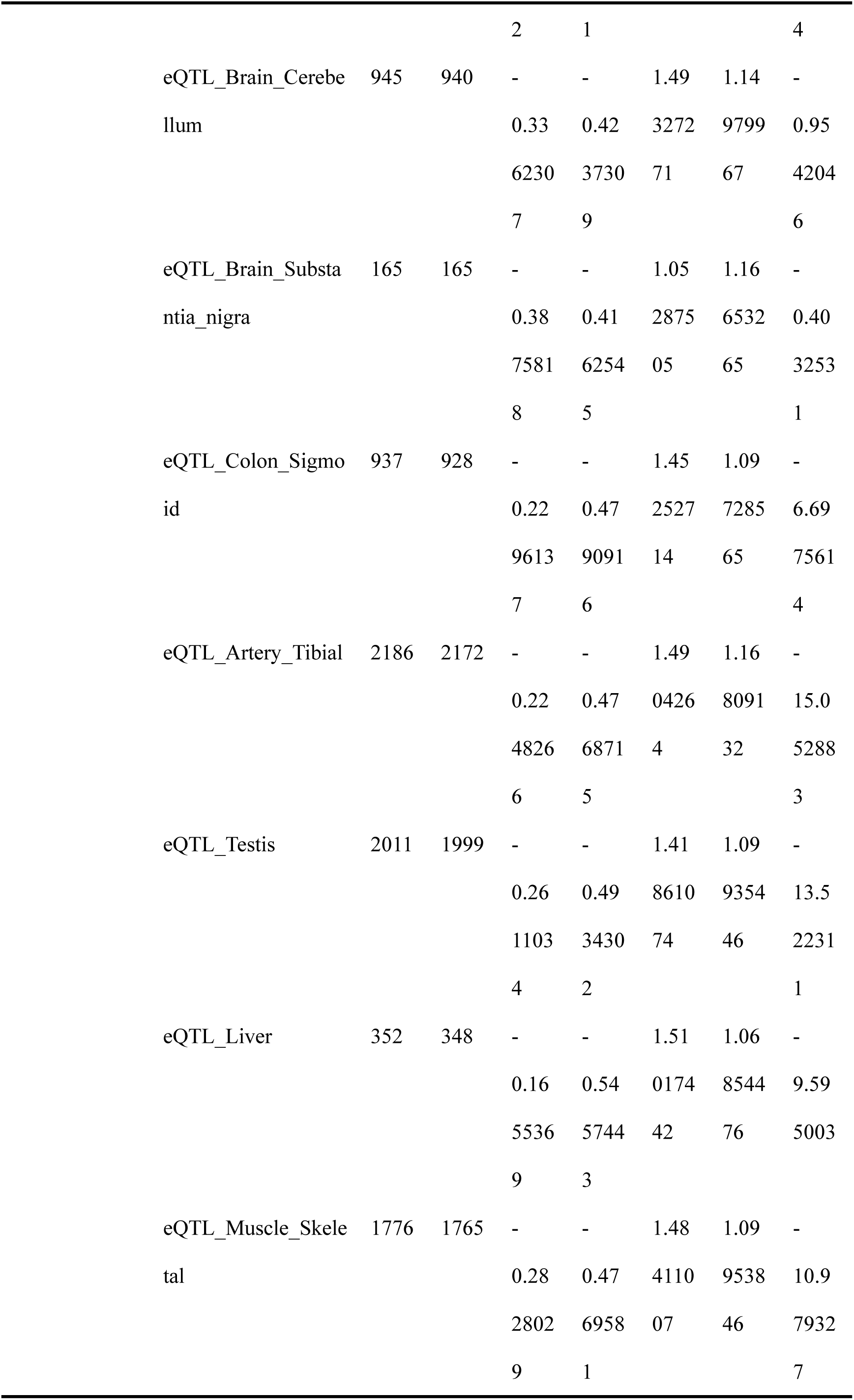
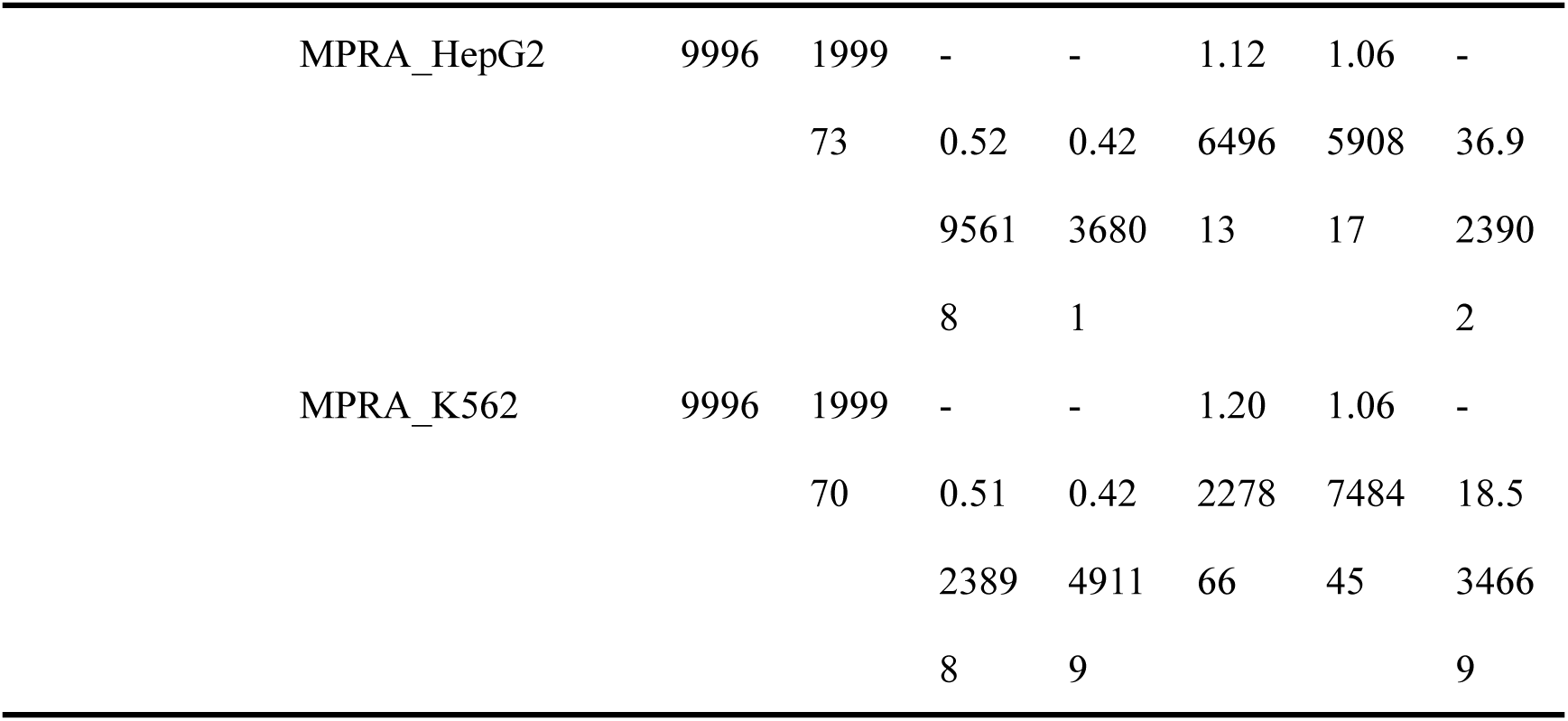
Distribution of Phylop-100way scores across variant effect prediction datasets.

**Supplementary Table 2.** Summary statistic of the long-range interaction dataset.

| Cell Type | Interaction | # Train Set | # Validation Set | # Test set |
| --- | --- | --- | --- | --- |
| tB | PPI | 141,254 | 15,695 | 5,442 |
|  | PEI | 153,664 | 17,074 | 5,358 |
| FoeT | PPI | 125,086 | 13,899 | 4,362 |
|  | PEI | 116,329 | 12,926 | 4,008 |
| Mon | PPI | 118,843 | 13,205 | 4,116 |
|  | PEI | 141,379 | 15,709 | 5,016 |
| nCD4 | PPI | 145,621 | 16,181 | 5,004 |
|  | PEI | 152,257 | 16,918 | 5,094 |
| tCD4 | PPI | 136,102 | 15,123 | 4,596 |
|  | PEI | 140,832 | 15,648 | 4,884 |
| tCD8 | PPI | 134,613 | 14,958 | 4,488 |
|  | PEI | 142,284 | 15,810 | 4,896 |

PPI: promoter-promoter interaction; PEI: promoter-enhancer interaction.

**Supplementary Table 3.** List of evaluated genomic language models.

| Model | Pretraining Data | Model Architecture | Tokenization Strategy | Pretraining Objective | Number of Parameters |
| --- | --- | --- | --- | --- | --- |
| Caduceus-ps<sup>2</sup> | Human | Mamba | Single- | MLM | 1.9M |

|  | genome |  | nucleotide |  |  |
| --- | --- | --- | --- | --- | --- |
| HyenaDNA-1M<sup>3</sup> | Human genome | Hyena | Single-nucleotide | CLM | 6.6M |
| DeepGene<sup>4</sup> | Human pan-genome | Transformer | BPE | MLM | 85M |
| GPN-MSA<sup>5</sup> | MSA from 100 vertebrates | Transformer | Single-nucleotide | MLM | 86M |
| DNABERT-3mer<sup>6</sup> | Human genome | Transformer | Overlapped 3-mer | MLM | 89M |
| DNABERT-2<sup>7</sup> | 135 species genomes | Transformer | BPE | MLM | 117M |
| OmniNA-220M<sup>8</sup> | 172 species genomes | Transformer (LLaMA) | BPE | CLM | 220M |
| LucaOne<sup>9</sup> | DNA, RNA, protein sequences from 169,861 species | Transformer | Single-nucleotide | MLM | 1.8B |
| NT-2.5B-MS<sup>10</sup> | 850 species genomes | Transformer | Non-overlapped 6-mer | MLM | 2.5B |
| GENERator-3B<sup>11</sup> | Multi-species genomes | Transformer | Non-overlapped 6-mer | CLM | 3B |
| Evo2-7B<sup>52</sup> | Genome sequences from 128M species | SSM (StripedHyena 2) | Single-nucleotide | CLM | 7B |

Note: Due to the design of the GPN-MSA model, it was only evaluated on the variant effect prediction application.

**Supplementary Table 4.**
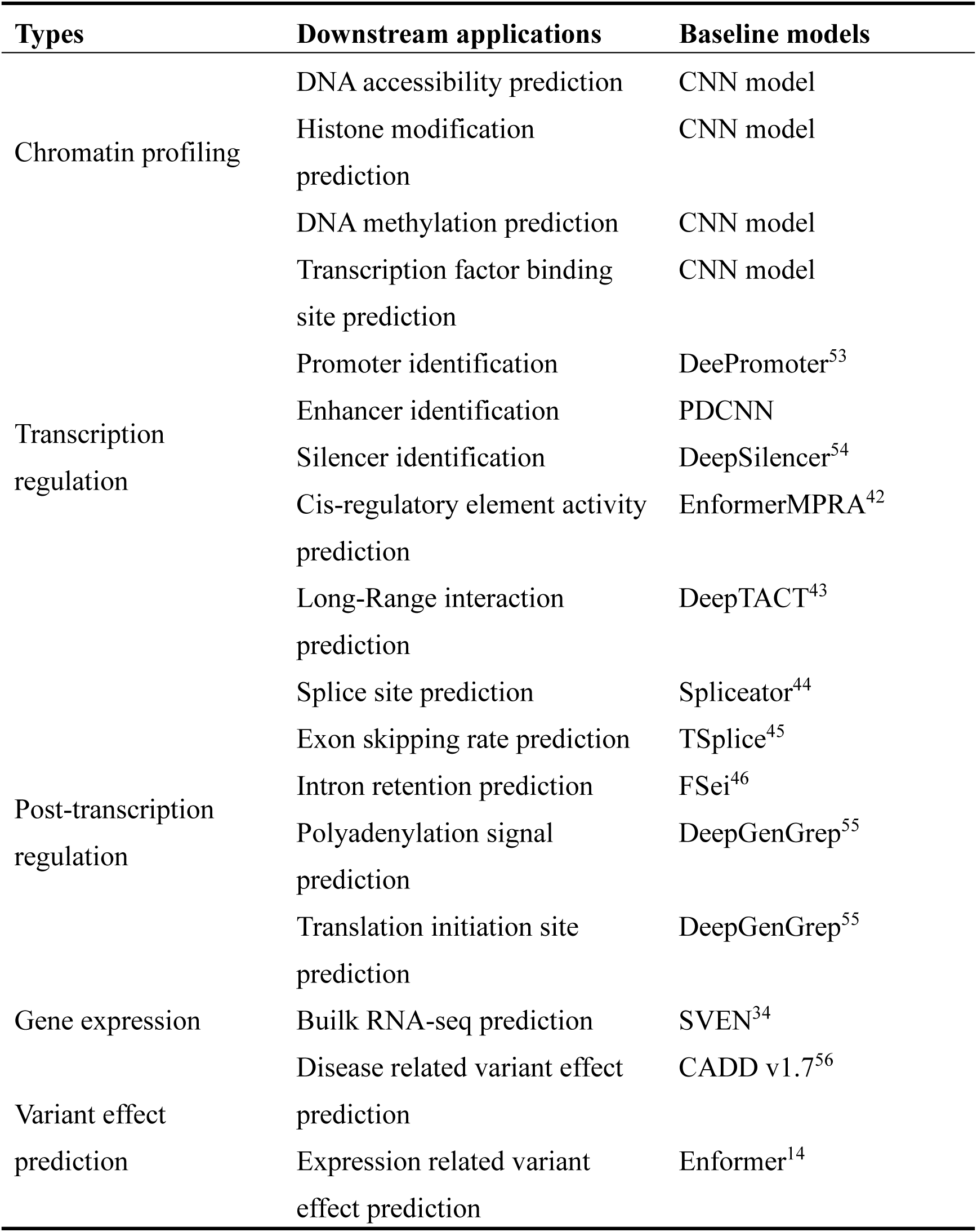
Non-gLM baselines for benchmarking.

**Supplementary Table 5.** Genomes for training multi-species gLM.

| Species | GenBank assembly |
| --- | --- |
| *Homo sapiens* | GCA_009914755.4 |
| *Pan troglodytes* | GCA_028858775.3 |
| *Macaca mulatta* | GCA_003339765.3 |
| *Canis lupus familiaris* | GCA_011100685.1 |
| *Mus musculus* | GCA_000001635.9 |
| *Rattus norvegicus* | GCA_036323735.1 |
| *Monodelphis domestica* | GCA_027887165.1 |

**Supplementary Table 6.**
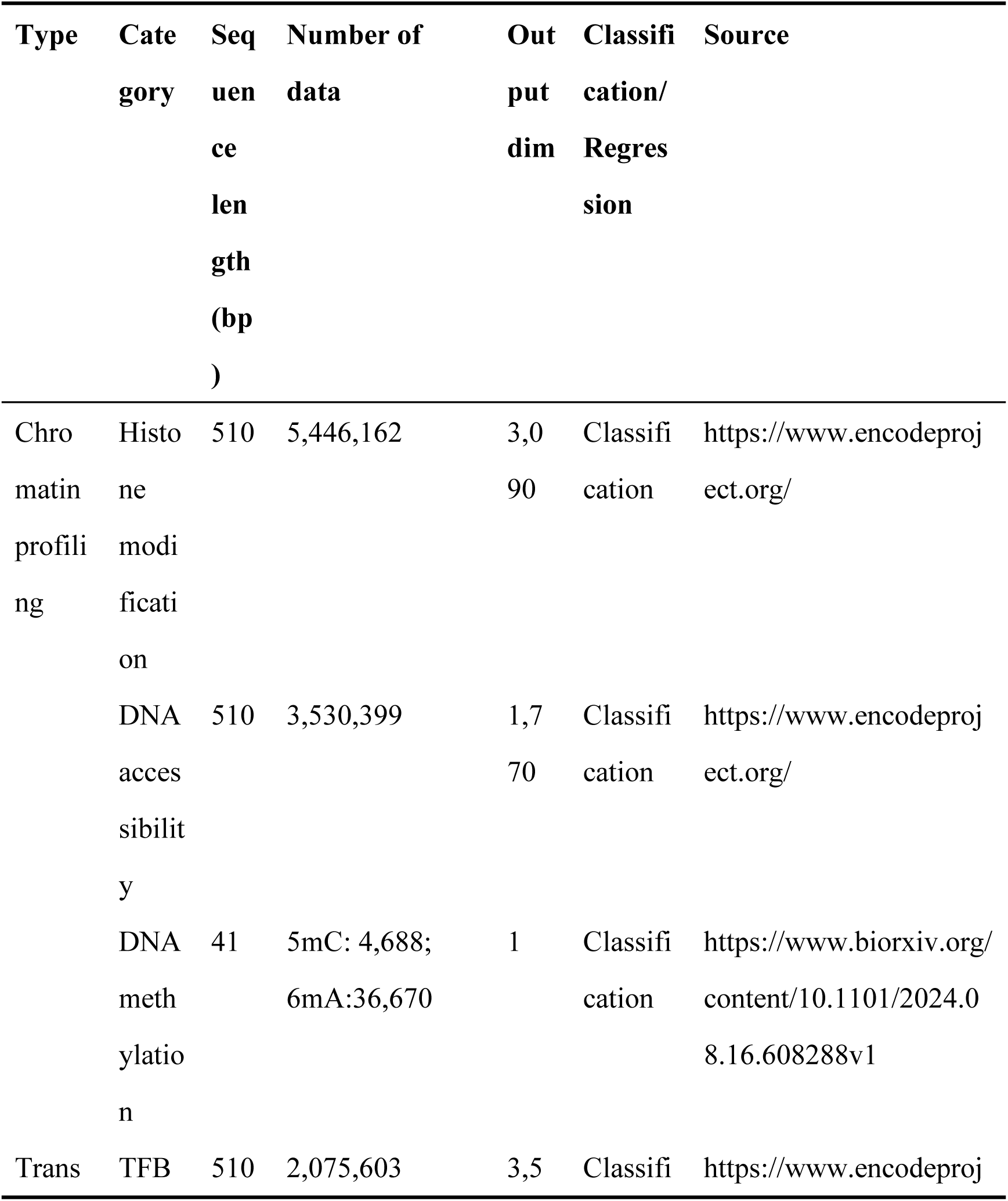
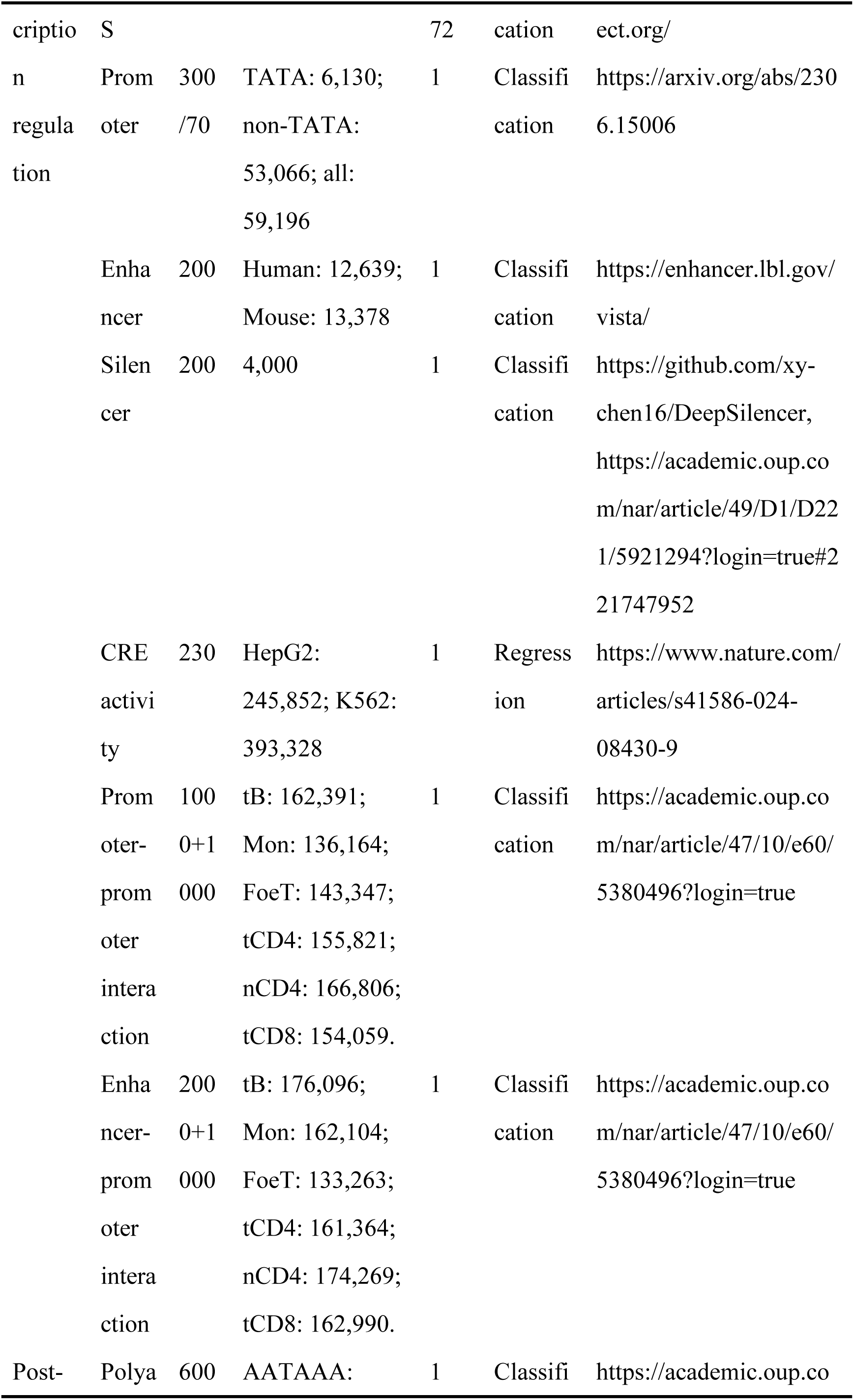
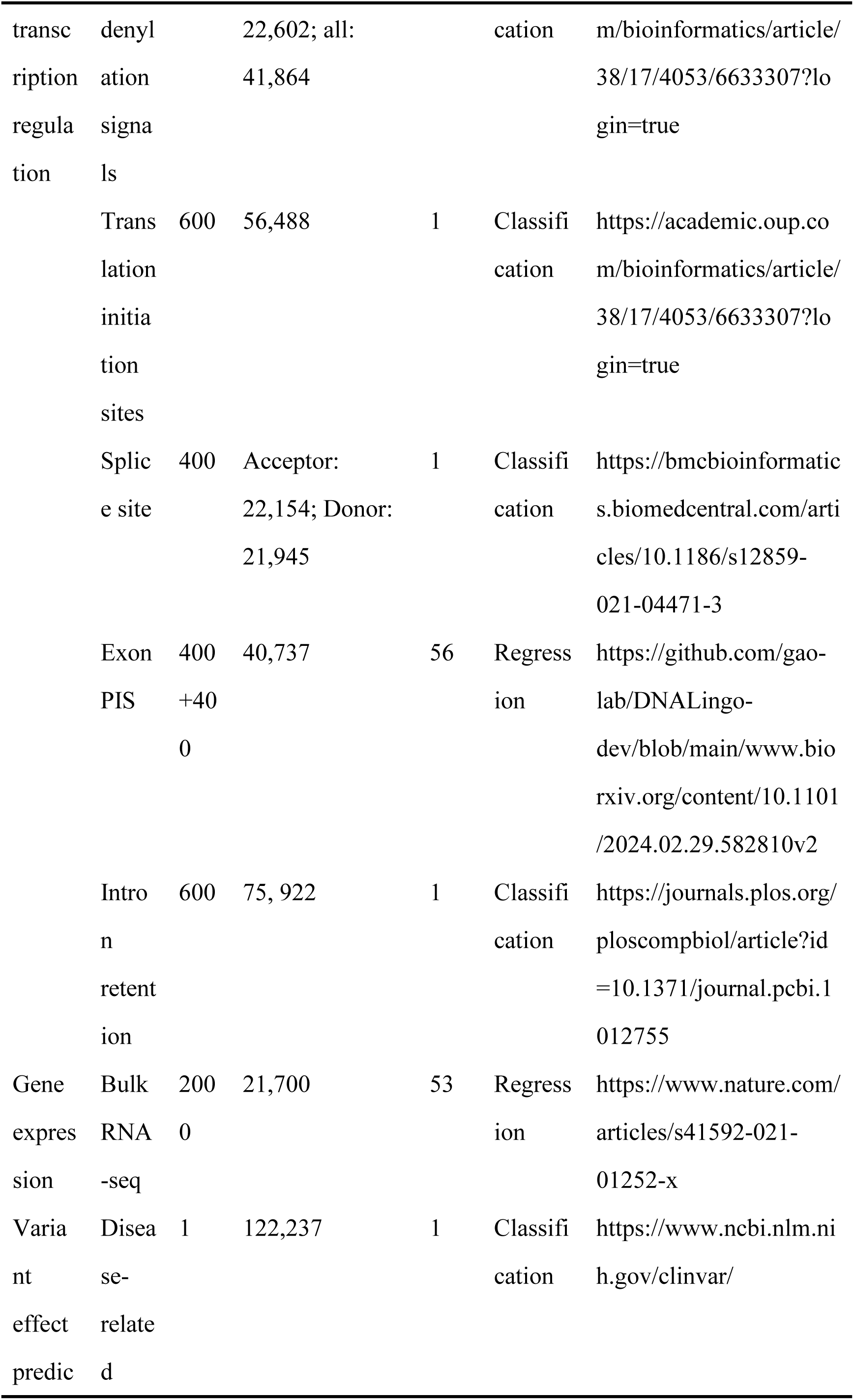
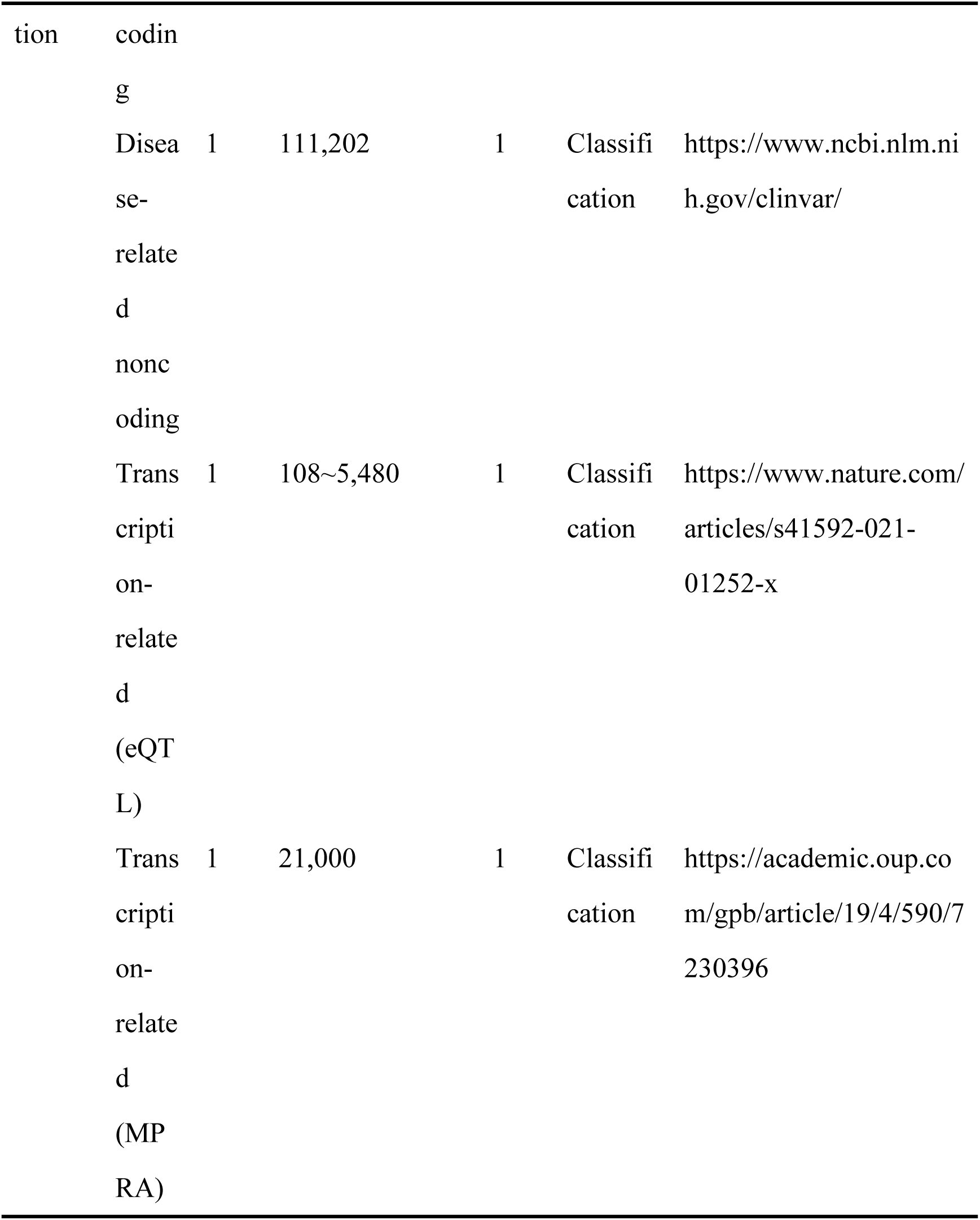
Complete list of the source datasets.

